# Contribution of epigenetic changes to escape from X-chromosome inactivation

**DOI:** 10.1101/2021.03.03.433635

**Authors:** Bradley P. Balaton, Carolyn J. Brown

## Abstract

**Background:** X-chromosome inactivation (XCI) is the epigenetic inactivation of one of two X chromosomes in XX eutherian mammals. The facultatively heterochromatic inactive X chromosome acquires many chromatin changes including DNA methylation and histone modifications. Despite these changes, some genes escape or variably escape from inactivation, and to the extent that they have been studied, epigenetic marks correlate with expression.

**Results:** We downloaded data from the International Human Epigenome Consortium and compared previous XCI status calls to DNA methylation, H3K4me1, H3K4me3, H3K9me3, H3K27ac, H3K27me3 and H3K36me3. At genes subject to XCI we found heterochromatic marks enriched, and euchromatic marks depleted on the inactive X when compared to the active X. Similar results were seen for genes escaping XCI although with diminished effect with H3K27me3 being most enriched. Using sample-specific XCI status calls made using allelic expression or DNA methylation we also compared differences between samples with opposite XCI statuses at variably escaping genes. We found some marks significantly differed with XCI status, but which marks were significant was not consistent between genes. We trained a model to predict XCI status from these epigenetic marks and obtained over 75% accuracy for genes escaping and over 90% for genes subject to XCI. This model allowed us to make novel XCI status calls for genes without allelic differences or CpG islands required for other XCI status calling methods. Using these calls to examine a domain of variably escaping genes, we saw XCI status vary at the level of individual genes and not at the domain level.

**Conclusion:** Here we show that epigenetic marks differ between genes that are escaping and those subject to XCI, and that genes escaping XCI still differ between the active and inactive Xs. We show epigenetic differences at variably escaping genes, between samples escaping and those subject to XCI. Lastly we show gene-level regulation of variably escaping genes within a domain.

## Introduction

In eutherian mammals, one of the two X chromosomes is epigenetically inactivated in XX females in order to achieve dosage compensation with XY males through a process known as X-chromosome inactivation (XCI) (see Balaton, 2018 for a review [1]). This inactivation is incomplete, as approximately 12% of genes consistently escape from XCI in humans [2], here defined as having at least 10% expression from the inactive X (Xi) as compared to the active X (Xa) [3]. There is growing interest in genes that escape XCI for their possible roles in sex-biased disease. Having two active copies of a gene offers additional protection from loss of function mutations linked to cancer [4] and likely underlies other sex-biased diseases and sex chromosome aneuploidies [5]. Genes that escape XCI tend to have sex-biased expression, being higher in males for genes that are also on the Y chromosome while being higher in females if the gene is only on the X [6]; genes with Y homology complicate this further by having different expression levels across tissues for the X and Y homologs [7]. However, relatively little is known about how a gene can be expressed from the midst of heterochromatin.

The pseudoautosomal regions (PAR) are located at the far ends of the X and maintain their ability to pair and recombine with the Y [8]. PAR1 contains approximately 30% of the genes described as escaping from XCI [2]. The short arm of the X chromosome near PAR1 is enriched in genes escaping from XCI, while the long arm, that contains XIST - the gene responsible for initiating XCI - is enriched in genes subject to XCI [9]. Genes escaping from XCI are often found clustered together, with some convergence with topologically associated domains (TADs) [10]. In addition to genes that consistently escape from XCI (sometimes called constitutive escape), a further 8% of genes have been found to vary their XCI status between different tissues or individuals (termed variable or facultative escape [2] (reviewed in [5]), and another 7% of genes were found to be discordant between the studies identifying them [2]. Variably escaping and discordant genes were found to be enriched at boundaries between clusters of genes with opposite XCI statuses [2]. The factors determining XCI status remain unresolved with the above evidence suggesting regional control, but there are also lone genes that escape XCI while flanked with genes subject to XCI [2] and even genes with two transcription start sites (TSSs) with opposite XCI status [11, 12].

Many methods have been used to identify which genes escape from XCI (reviewed in [13]). The gold-standard approach is to compare expression between the Xi and Xa within a sample [6, 9, 14], which requires a heterozygous SNP within an exon to differentiate Xi from Xa expression. Such expressed SNPs can be rare, and additionally, which X chromosome is inactivated is normally random throughout a subject’s body, precluding assigning an allele to the Xi. Some samples are naturally skewed so that the same X is inactivated in >90% of their cells, which allows allelic analysis. The frequency of such skewing is increased in blood [15], with age [16] and also in cancer, which generally arises clonally [17]. Cells that have become monoclonal during cell culture and those skewed due to deleterious alleles on one copy of the X chromosome have also been used for allelic expression analysis [3]. Single cell RNA-sequencing (RNA-seq) has also been used to avoid the need for clonal cell populations when using Xi/Xa expression to make XCI status calls [18]. Single cell RNA-seq can additionally see variation in XCI status between different Xi alleles within the same sample and heterogeneity has even been observed in XCI status of a gene in cells with the same Xi [6, 19].

Beyond the direct examination of allelic expression, the modifications to DNA and chromatin that accompany XCI can be used as surrogates to determine if a gene is inactivated. For these features it is unclear if the mark enables or reflects XCI status. Promoter DNA methylation (DNAme) at CpG islands is an epigenetic mark that is strongly predictive of a gene’s XCI status and has been used to differentiate genes that escape XCI from those subject to XCI without the need for heterozygous SNPs or skewed Xi choice [20]. Low promoter DNAme on the Xa, as evidenced by low DNAme in male samples, is necessary to allow detection of DNAme differences on the Xi. Genes that escape from XCI will also have low DNAme in females, with both the Xi and Xa unmethylated, while genes that are subject to XCI will have intermediate DNAme, with the Xa unmethylated and the Xi methylated. Other epigenetic marks such as histone marks have been reported to be correlated with a gene’s XCI status. Active marks such as H3K4me2/3, H3K9ac, H3K27ac, H3K9me1, RNA polymerase II and transposase accessibility are enriched at genes escaping from XCI, while inactive marks such as H3K9me3, H4K20me3, H3K27me3 and macroH2A are enriched at genes subject to XCI [13, 21, 22], reviewed in [13]. A predictive model using many epigenetic and genetic features in mice was able to predict a gene’s XCI status accurately 78% of the time [23] and in humans a model obtained over 80% accuracy using only genomic repeats [24].

In order to understand how genes are (or are not) silenced on the Xi, in a chromosome-wide fashion, we take advantage of the growing genome-wide epigenomic datasets to correlate XCI status and multiple epigenetic marks. The strength of these correlations permitted the development of an epigenetic predictor of XCI status, allowing prediction of escape. Within a single region we observed control of silencing/escape at the single gene level.

## Results

To understand the interplay of epigenetic marks and XCI status we sought a dataset with both a broad range of epigenetic marks and matched expression data to determine XCI status. We thus turned to data from the International Human Epigenome Consortium (IHEC), which has standardized chromatin immunoprecipitation sequencing (ChIP-seq) for the core histone marks H3K4me1, H3K4me3, H3K9me3, H3K27ac, H3K27me3 and H3K36me3 along with whole genome bisulfite sequencing (WGBS) to examine DNAme. We specifically used data from the Center for Epigenome Mapping Technologies (CEMT) as these samples were derived from cancer and thus were anticipated to have a high frequency of skewed XCI, allowing us to use allelic expression to determine XCI status [12]. As cancer is known to have epigenetic changes, we additionally examined data from Core Research for Evolutional Science and Technology (CREST), another group within IHEC, thus allowing us to determine whether any trends that we observed in the CEMT data were due to the samples being cancer-derived. However, the CREST samples had less sequencing depth, fewer females (only nine), and could only be examined for DNAme and histone marks. Samples are listed in Table S1.

### Histone marks differ with sex and XCI status

We first compared the levels of histone modifications within 500bp upstream of a gene’s TSS (except for the mark H3K36me3 that is associated with gene bodies and so was examined at exons [25]) with sex and published XCI status calls derived from a synthesis of various approaches (hereafter referred to as meta-status) [2]. For the purposes of this and subsequent analyses, genes in the PAR were not included with genes escaping from XCI as they may be epigenetically distinct, especially when comparisons with males are included.

We found that most marks had a significant difference (p-value < 0.01) for the median level per transcript between males and females, at genes escaping and subject to XCI in both datasets (**Table 1**). Fewer marks showed significant differences between genes escaping XCI and those subject to XCI within each sex. The euchromatic marks (H3K4me1, H3K4me3, H3K27ac, and H3K36me3) were significantly different between transcripts subject to XCI and those escaping from XCI in both CEMT and CREST females, while the heterochromatic marks (H3K9me3, H3K27me3 and DNAme) were only significantly different within the CREST dataset. Comparing

**Table 1:** Significance of differences in epigenetic marks between sex and transcripts of opposing XCI status. Values shown are Benjamini-Hochberg adjusted p-values. Those with an adjusted p-value<0.01 are shown in bold. CEMT analysis is shown, see Table S2 for analysis of CREST data and other comparisons. There are 102 transcripts escaping XCI and 993 transcripts subject to XCI included in these analyses.

|  | Escape XCI,<br>Male vs Female | Subject to XCI,<br>Male vs Female | Within females,<br>escape vs subject | Within males,<br>escape vs subject |
| --- | --- | --- | --- | --- |
| H3K4me1 | **4.56E-09** | **1.99E-09** | 1.12E-02 | **7.29E-05** |
| H3K4me3 | **4.54E-07** | **4.54E-07** | **6.29E-06** | 9.48E-01 |
| H3K9me3 | **5.24E-06** | **2.24E-170** | 1.12E-02 | **4.79E-03** |
| H3K27ac | **7.95E-04** | **7.29E-05** | **6.05E-06** | **3.63E-03** |
| H3K27me3 | **3.78E-07** | **5.24E-276** | 9.48E-02 | 2.61E-01 |
| H3K36me3 | 1.12E-02 | **4.56E-09** | 6.78E-01 | 9.48E-01 |
| DNAme | 9.48E-01 | **1.48E-161** | **2.75E-18** | 7.50E-01 |
| expression | 7.37E-01 | 2.57E-01 | 5.23E-01 | 5.93E-01 |

XCI statuses within males gave the fewest significantly different marks, as was expected. We tested the male:female differences at each transcript for each mark and observed two blocks of correlations (**Figure 1a**). The heterochromatic marks were moderately correlated together while the euchromatic marks had a weaker correlation. Expression was most correlated to H3K36me3, and its correlation to the other euchromatic marks was as low as its correlation with the heterochromatic marks.

**Figure 1:**
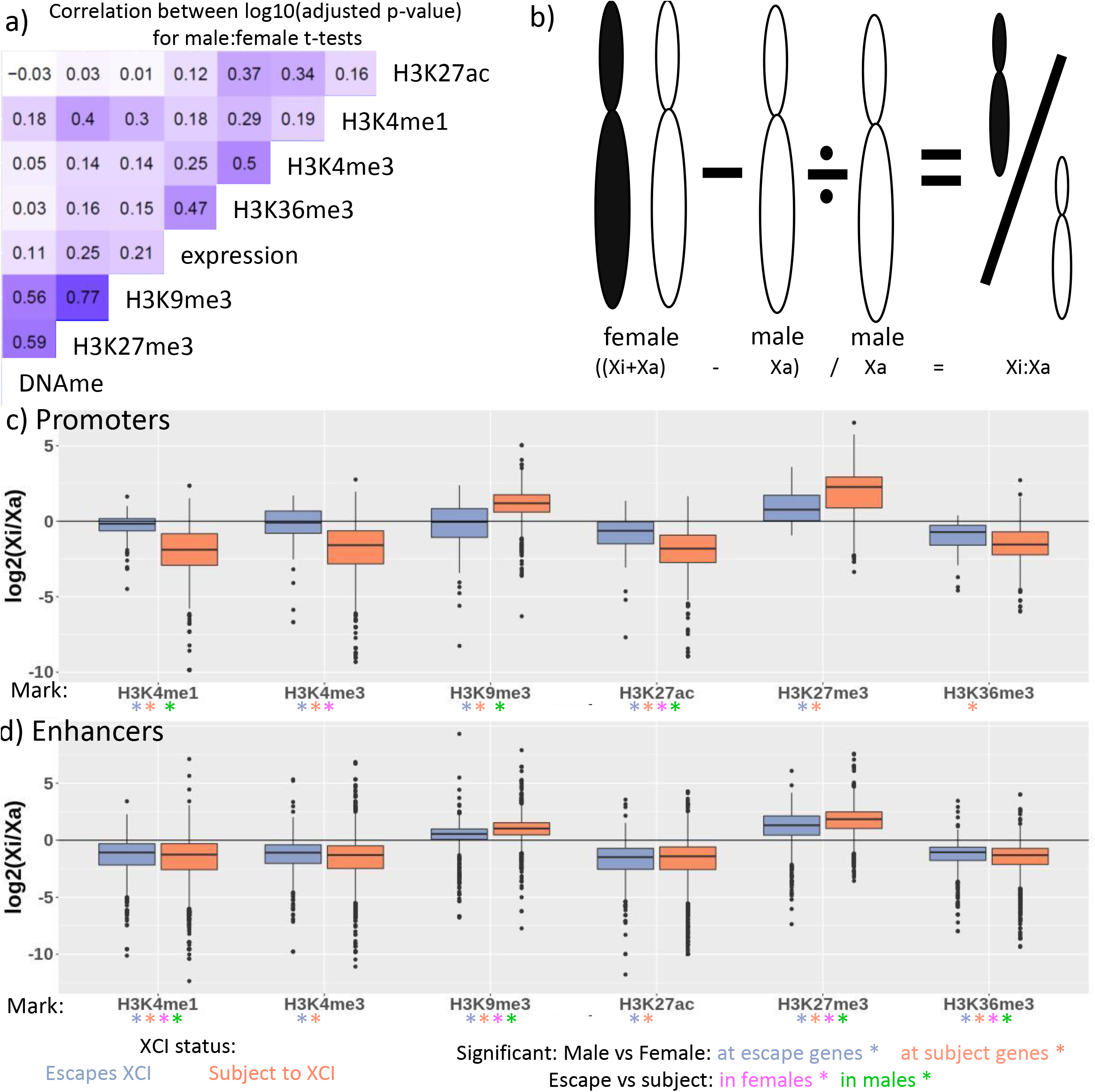
The Xi has more heterochromatic and less euchromatic marks than the Xa. (a) Correlations between log_10_(adjusted p-values) for t-tests between male and female samples per transcript for each mark. (b) a diagram showing our Xi/Xa calculation. (c-d) Log_2_(Xi/Xa ratio) for the histone marks examined here at promoters (c) and enhancers (d), split by XCI meta-status. Data from CEMT is shown. Significance for the various t-tests featured in Table S2 are shown by the differently colored star.

To visualize the differences between the Xi and Xa, we calculated the Xi to Xa fold change for each mark by taking log2 of the female:male difference (the contribution from the Xi) and dividing it by the male value (the contribution from the Xa) (**Figure 1b-c, Figure S1** for CREST**, Table S2** for values). Heterochromatic marks are generally higher on the Xi than Xa, especially for transcripts subject to XCI. H3K27me3 has a higher Xi:Xa fold change than H3K9me3 in both XCI statuses and both marks are highest at transcripts subject to XCI. For euchromatic marks, the Xi:Xa ratio is close to 1:1 at transcripts escaping XCI, and lower for transcripts subject to XCI. H3K36me3 is reduced on the Xi for both XCI statuses with the Xi being approximately one half of the Xa. For transcripts subject to XCI, the healthy CREST samples had less of an Xi to Xa difference than the CEMT cancer samples. At transcripts escaping from XCI the differences were weaker, and more variable between the datasets. There are almost ten times as many transcripts subject to XCI as there are escaping XCI, which partially explains the stronger p-values at transcripts subject to XCI.

H3K27me3 showed the largest change between the Xi and Xa, yet was not significantly different between XCI status in females (or males). We thus examined the proportion of transcripts that showed significant male:female differences, subdivided by XCI metastatus (**Table S3**). While DNAme and H3K9me3 show substantial enrichment in females predominantly at transcripts subject to XCI, we found H3K27me3 significant in over 85% of the transcripts in any XCI status. To validate this broad enrichment across the Xi we downloaded H3K27me3 data from ENCODE and found a similar trend to the CEMT data, with over 70% of transcripts in the escaping, subject to XCI and variably escaping categories being significantly different between the sexes. We analyzed chromosome 7 as an example autosome and saw a much lower percentage of transcripts with significant male-female differences for H3K9me3 and H3K27me3. This proportion was lower than even transcripts escaping from XCI, so even transcripts that escape from XCI have a significant increase of these heterochromatic marks in females relative to males.

Given the sex differences across the X chromosome we wished to examine how the marks differed across sex and XCI status by generating meta-gene plots extending 50kb up and downstream of genes escaping or subject to XCI, in females and males (**Figure S2**). H3K4me3 and H3K27ac look similar between XCI statuses and sexes, except for having a higher peak at the TSS for genes escaping XCI in females. H3K9me3 and H3K27me3 peak at the TSSs for genes subject to XCI in females, with a smaller peak at genes escaping XCI. We also see these heterochromatic marks lower in the gene bodies of genes escaping XCI than in the 50kb up and downstream of the gene. In H3K27me3, the regions up and downstream of genes escaping from XCI are actually higher near genes escaping XCI than at those subject to XCI in both females and males. H3K36me3 looks similar between males and females, and is higher at genes escaping XCI than at genes subject to XCI. For all marks, the standard deviation across genes with each XCI status was large, calling into question whether the differences would be predictive for individual genes, as has been found for DNAme (see **Table S2**). DNAme at promoters was significantly different between males and females at genes subject to XCI and in females between genes subject to XCI and those escaping from XCI. Expression was not found significantly different in any of the comparisons here.

In addition to our promoter and gene-based analysis, we also compared histone marks at enhancers annotated to genes on the X [26] and found that all marks showed significant, although small, differences between males and females,for both XCI statuses (**Figure 1d, Table S2** for values). We further considered whether the enhancer was found within the gene to ensure that differences were not arising simply due to expression of the gene altering chromatin; however, most marks remained significant regardless of location. At enhancers, heterochromatic marks, along with H3K4me1 and H3K36me3, differ significantly with XCI status of their associated gene, in both sexes. Looking at the Xi:Xa fold change at enhancers (**Figure 1d,** see **S3** for division into genic and intergenic enhancers), all of the heterochromatic marks were higher on the Xi than the Xa, while euchromatic marks were higher on the Xa than the Xi. This did not differ greatly between enhancers that were annotated to interact with genes escaping and genes subject to XCI.

### Epigenetic marks correlate with sample-specific changes in XCI status

Heterogeneity of XCI status means that our use of meta-status will be less accurate than individual-specific XCI calls. We thus used our Xi/Xa expression-based XCI status calls in a subset of the CEMT samples (as reported previously [12]) to analyze the interaction between XCI status and epigenetic marks, made within the same sample. For this sample-specific expression analysis we could only examine females with skewed XCI such that the same X was inactive in all cells. Across the eight skewed samples, 30 genes escaped XCI, 202 genes were subject to XCI and 8 genes variably escaped from XCI (requiring at least two samples with each XCI status to be called variable) (**Figure 2a**, **Table S4** for XCI status calls)). Two of the genes found to variably escape herein were previously called as variably escaping from XCI, while five were previously designated escaping XCI and one subject to XCI.

**Figure 2:**
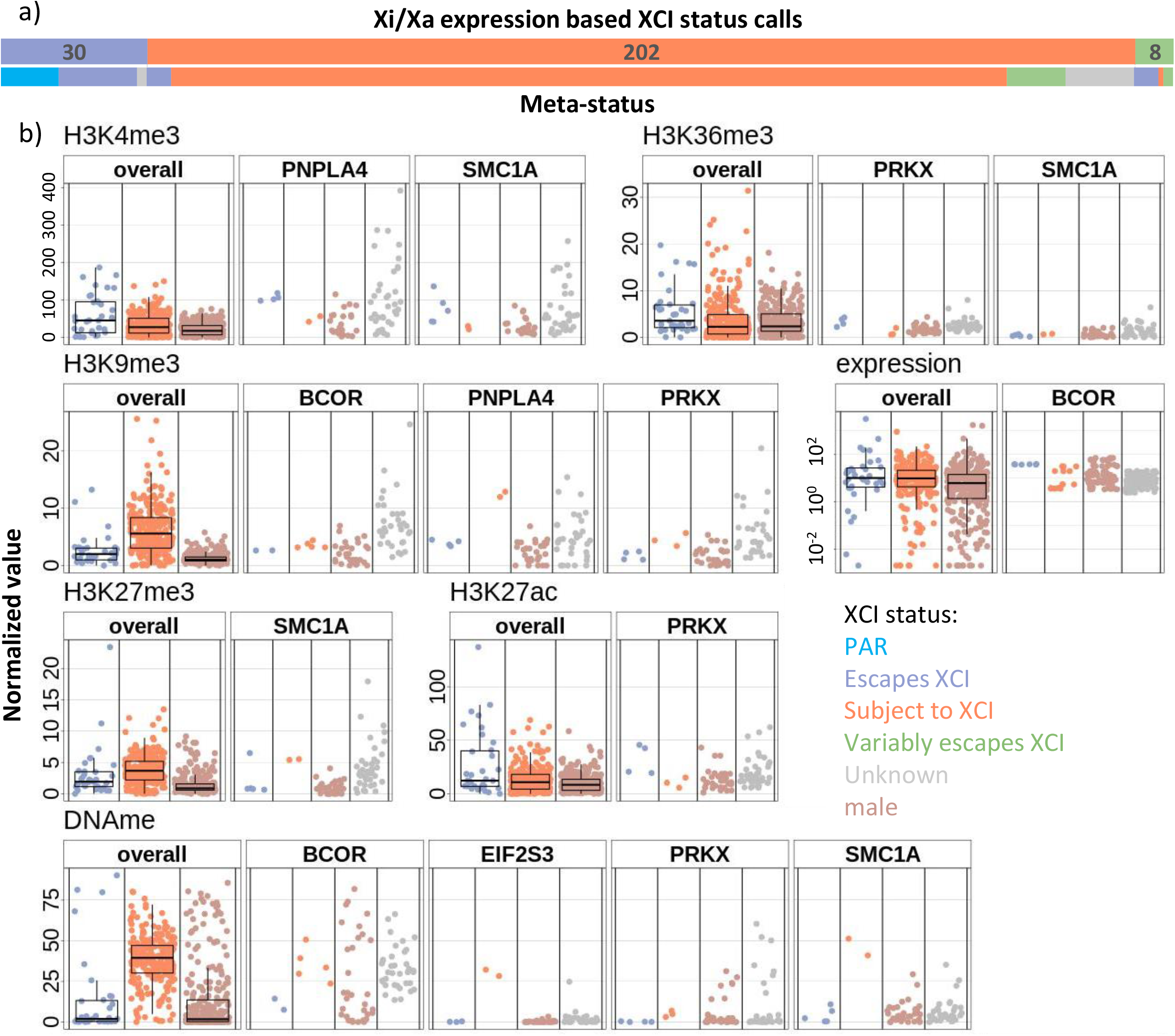
Epigenetic marks do not change consistently with XCI status for variably escaping genes. (a) The number of genes with each XCI status call across all samples, by Xi/Xa expression with their distribution by meta-status underneath. (b) The interaction of histone marks and Xi/Xa expression determined XCI status. On the left for each mark is a comparison to the overall XCI status across samples per gene and on the right are shown the variably escaping genes that had significant differences in the histone mark between samples that were subject to or escaping from XCI. A p-value of 0.05 was used for significance. Unknown XCI status is for samples that were uninformative in the Xi/Xa expression analysis. Expression is on a log_10_ scale while the others are on a linear scale.

The sample-specific XCI status calls determined by allelic expression are more precise; however their use did not substantively alter the histone mark representations found using meta-status calls (**Table S2**). The histone marks that were found significant in all three comparisons (CEMT marks vs meta-status, CREST marks vs meta-status, CEMT marks vs CEMT Xi/Xa calls) are H3K4me3 and H3K9me3 (in all but the male XCI comparison), H3K27me3 (between sexes for both XCI statuses), H3K27ac and DNAme (between sexes at genes subject to XCI and between XCI statuses in females), and H3K4me1 and H3K36me3 (only between sexes at genes subject to XCI). Expression was not different between sexes or XCI statuses here. An additional benefit of having both histone marks and XCI status on individual samples is the ability to examine how histone marks correlate with XCI status at variably escaping genes.

Genes that variably escape from XCI provide a unique opportunity to study differences between genes escaping vs subject to XCI in the same genomic context. All of the marks available except for H3K4me1 were significantly different (p-value <0.05) between samples escaping XCI vs those subject to XCI in at least one of the eight variably escaping genes, but never for the majority of genes (**Figure 2b, Table S5**). Consistent with the associations seen for genes subject to or escaping from XCI, when active marks were significantly different, they tended to be higher in samples escaping XCI, while inactive marks were lower in samples escaping XCI (**Table S6**). The exception to this is H3K36me3 in gene bodies. For one gene the samples subject to XCI had higher H3K36me3, while in another gene it was the samples escaping XCI that were higher.

DNAme was the most consistent mark differentiating samples escaping from those subject to XCI, being seen significantly different in four out of the eight variably escaping genes. The samples subject to XCI in *PRKX* had significantly higher DNAme, but were not above the DNAme thresholds for XCI status calls that we established previously [12]. The other three genes with significant DNAme differences showed a clear switch from a DNAme pattern matching genes escaping XCI to a pattern matching genes subject to XCI. *TIMP1*, one of the four genes that was not significant, has low CpG density and high male DNAme so was not expected to differ with XCI status. For the other three genes, the limited informative samples reduced the power to detect differences, although they may have had incorrect XCI status calls or there may be more complicated epigenetic processes involved. Interestingly, the two genes found to be variably escaping by both Xi/Xa expression and meta-status (*MED14* and *TIMP1*) did not show DNAme differences while many of the genes without meta-status calls of variable escape had significant DNAme differences. Thus we have confidence that these other genes (*BCOR, EIF2S3, PRKX* and *SMC1A*) are truly variable across these samples. Three of the variably escaping genes did not show significant differences at any of the examined marks; increasing the sample size might give us the power to see more consistent differences across variably escaping genes as some of these genes only had 2 informative samples per XCI status.

We considered a series of other potential contributors to variability in escape from XCI including differences in sequence, expression, or tissue. Testing whether the Xi allele or the genotype of nearby polymorphisms had any correlation with XCI status would require many more samples to allow for multiple testing correction, as we found an average of only six informative samples per gene. Two genes showed significant expression differences between samples that escaped XCI versus those subject to XCI (**Figure S4**). In *BCOR*, samples escaping XCI had higher expression across all exons, while in *EIF2S3* some exons were higher in samples subject to XCI while other exons were higher in samples escaping XCI. XCI status and expression per exon may be linked by different TSSs having different XCI status or possibly different tissues having different XCI status and dominant splicing variants. To test whether variable escape may be tissue-specific, XCI status per sample was compared with tissue of origin; only one of the eight genes showed tissue-specificity, *EIF2S3*. However, with only eight samples in three tissue types and being limited by heterozygous polymorphisms, there are likely other tissue-specific variable escape genes that were not identified here as many genes did not have multiple informative samples per tissue.

### Expanding sample-specific XCI status by using DNA methylation

To increase our sample size, we used promoter DNAme levels to determine XCI status across all genes within the larger 45 sample CEMT dataset, regardless of skewed XCI. Only TSSs with high CpG density and low male methylation were considered informative, and within this group we found 47 genes escaping XCI, 393 subject to XCI and 18 variably escaping across samples (**Figure 3a**, **Table S4** for XCI status calls). Our DNAme based calls had strong concordance with meta-status; there were no genes called as escaping XCI here that were previously called as subject to XCI, while only one of the genes called as subject to XCI here was previously called as escaping XCI. We included genes in the variably escaping from XCI category if at least one of their TSSs had 33% or more of its samples escaping XCI and another 33% or more samples subject to XCI. Additionally, one gene had opposing XCI statuses at separate TSSs and 36 had opposite XCI statuses across tissues (examples of genes with these variable escape scenarios are shown in **Figure 3b**). An additional 67 genes were found variably escaping in at least one tissue but were not identified as variably escaping from XCI in the larger dataset. Only *BCOR* was found variably escaping from XCI in the Xi/Xa expression-based calls and also found variably escaping here. In addition 96% of genes escaping and 87% of genes subject to XCI identified by Xi/Xa expression in these samples had concordant status in our DNAme based calls, with most of the discrepancies between calls being due to genes being called as variably escaping in only one of the datasets.

**Figure 3:**
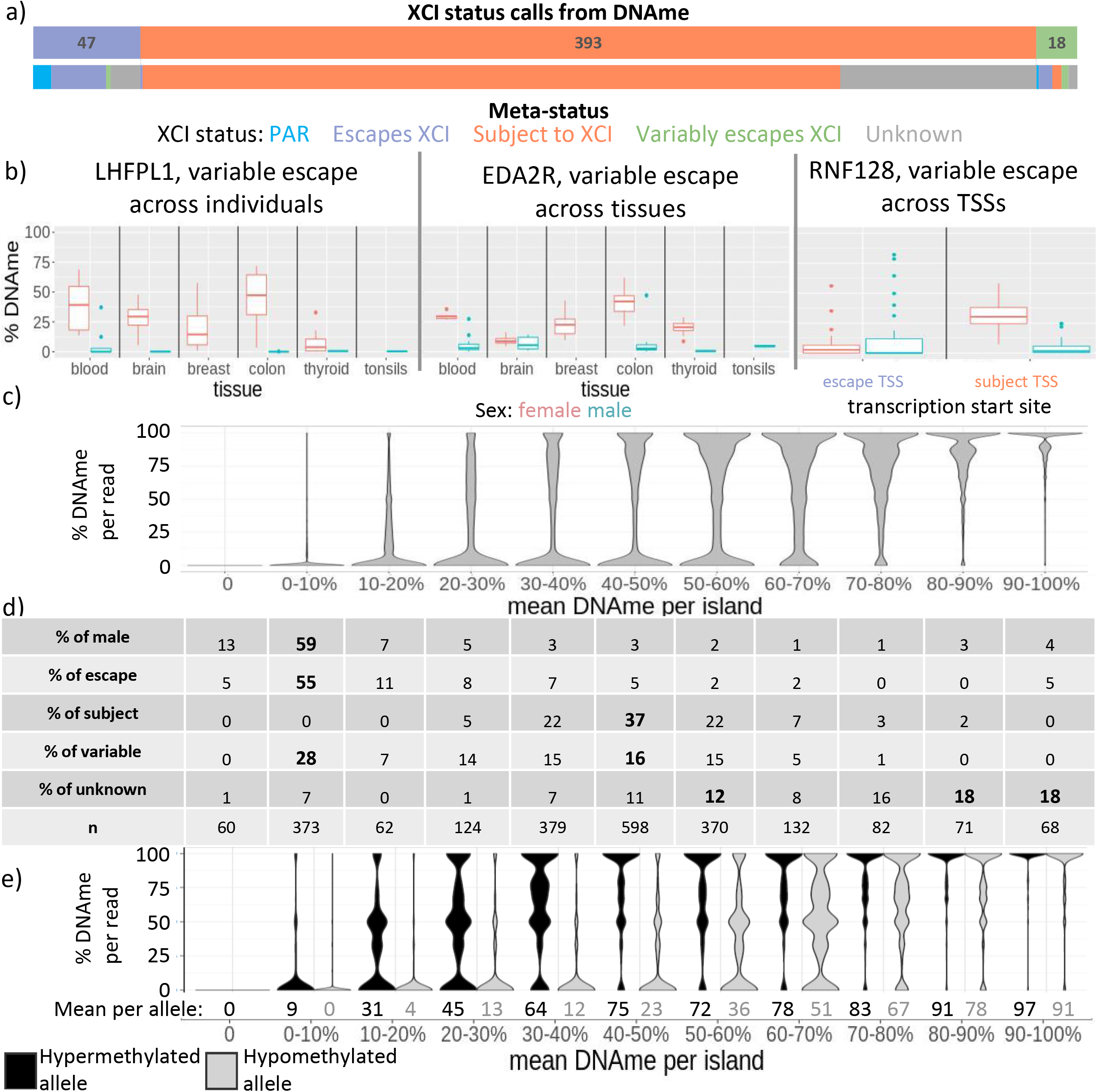
DNAme varies at genes variably escaping from XCI. (a) The number of genes with each XCI status call by DNAme, with their distribution by meta-status underneath. (b) From left to right: An example of a gene that variably escapes XCI across individuals (and within multiple tissues), a gene that variably escapes from XCI between tissues, and a gene that variably escapes from XCI between TSSs. (c) The percent DNAme per read for genes, binned together by their mean DNAme across the CpG island. Only reads overlapping the CpG island were included here. (d)The distribution of genes with each XCI status across the bins of mean DNAme per island. (e) Allelic DNAme, shown as the percent DNAme per read by allele. The mean DNAme across all reads per allele in each bin is shown underneath.

Comparing epigenetic marks to DNAme based XCI status calls, H3K4me1 and H3K36me3 were not significantly different between genes with opposite XCI status calls but the rest of the marks (H3K4me3, H3K9me3, H3K27me3 and H3K27ac) were very significant, with increased prominence of H3K9me3 (**Table S7**). We again compared epigenetic marks at variably escaping genes to see if they differed between samples in which the gene escaped XCI vs those in which it was subject to XCI. We categorized variable escape genes as those variably escaping across the dataset, across TSSs, across tissues or within specific tissues. For variable escape from XCI between individuals across the dataset, every mark examined was found to be significant (p-value<0.01) in at least one gene; however across all categories of variable escape from XCI, only DNAme, expression and H3K4me3 were significant in more than 25% of genes in any type of variable escape category (**Table 2)**. The direction of histone marks changes was less consistent than for Xi/Xa expression based XCI status calls, with the majority of genes still having higher active marks in genes escaping XCI and higher inactive marks in genes subject to XCI, but with many genes showing the opposite results (**Figure S5**).

**Table 2:**
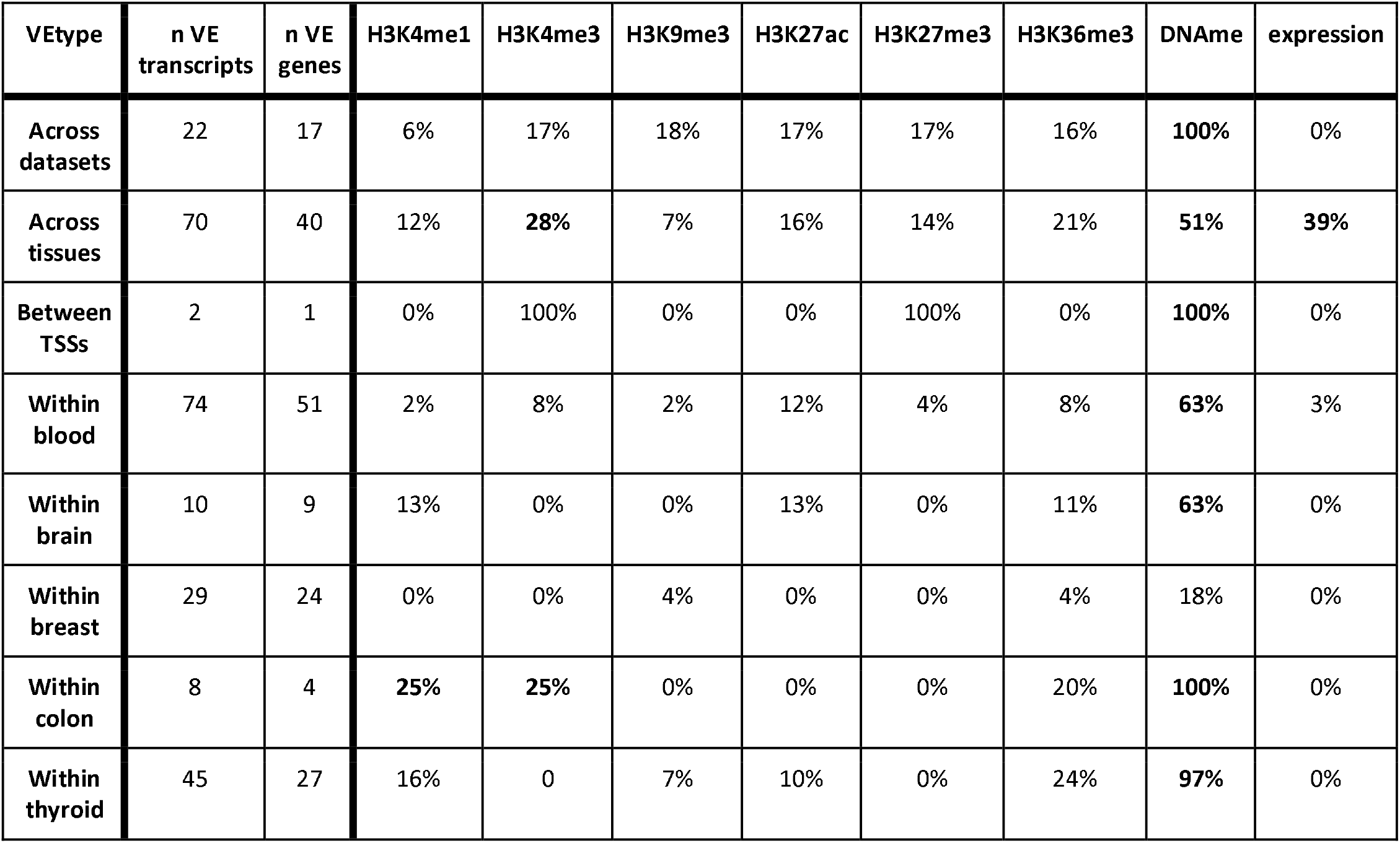
The percentage of variably escaping genes found by DNAme that have significant differences in epigenetic marks (BH corrected p-value<0.01). Different categories of variable escape are included on the left. The number of variably escaping (VE) transcripts found per category and the number of unique genes are also included. Categories with 25% or more variably escaping genes found significant are bolded, excluding variable escape between TSSs that only had 1 gene available.

| VType | n VE transcripts | n VE genes | H3K4me1 | H3K4me3 | H3K9me3 | H3K27ac | H3K27me3 | H3K36me3 | DNAm | expression |
| --- | --- | --- | --- | --- | --- | --- | --- | --- | --- | --- |
| Across datasets | 22 | 17 | 6% | 17% | 18% | 17% | 17% | 16% | **100%** | 0% |
| Across tissues | 70 | 40 | 12% | **28%** | 7% | 16% | 14% | 21% | **51%** | **39%** |
| Between TSSs | 2 | 1 | 0% | 100% | 0% | 0% | 100% | 0% | **100%** | 0% |
| Within blood | 74 | 51 | 2% | 8% | 2% | 12% | 4% | 8% | **63%** | 3% |
| Within brain | 10 | 9 | 13% | 0% | 0% | 13% | 0% | 11% | **63%** | 0% |
| Within breast | 29 | 24 | 0% | 0% | 4% | 0% | 0% | 4% | 18% | 0% |
| Within colon | 8 | 4 | **25%** | **25%** | 0% | 0% | 0% | 20% | **100%** | 0% |
| Within thyroid | 45 | 27 | 16% | 0 | 7% | 10% | 0% | 24% | **97%** | 0% |

We have previously seen that the average DNAme at genes subject to XCI was 38%, less than expected if the Xi were completely methylated, and that some genes subject to XCI had DNAme as low as 15% [12]. It seems likely that the lower female methylation reflects lower Xi methylation, but we further questioned whether lower XI methylation could predispose genes towards variable escape from XCI. Therefore, we examined the DNAme per WGBS read at CpG islands for the six female samples for which we had WGBS aligned reads, along with one male as a control. The reads for each gene in each sample were subdivided into 10% DNAme bins using the mean DNAme for the gene (**Figure 3c**). The genes in the 30-40% DNAme and 40-50% DNAme bins had a surprisingly low number of reads with high DNAme (24% and 35% of reads over 75% DNAme, respectively) so it appears to be that the majority of cells are partially methylated and not that some cells are methylated while others are not (**Table S8**). Over 80% of genes with a meta-status of subject to XCI had a mean DNAme between 30% and 60%, while over 60% of genes with a meta-status of escape from XCI and 70% of males across all categories had mean DNAme less than 10% (**Figure 3d**). Variably escaping genes were found distributed in the range where genes escaping and subject to XCI were found; however, genes with intermediate 20-30% DNAme had more variably escaping genes than genes with a consistent XCI status. Genes with no known XCI status tended to have high DNAme, with over half of them having 70% DNAme or higher. Intermediate DNAme (reads with 33-66%) is found most frequently in the 20-30% and 70-80% bins.

While the bimodal appearance of the DNAme reads reflects that the Xi and Xa are behaving differently, the intermediate reads could be derived from either. To differentiate DNAme from the Xa and Xi, we examined DNAme per read overlapping heterozygous SNPs within 2kb of TSSs. In addition to the usual limitations of mapping allelic reads, we had to exclude C<>T and G<>A polymorphisms as the bisulfite conversion step in WGBS converts unmethylated C to T and on the opposite strand this appears as a G to A conversion. Separating genes into the same 10% bins of mean DNAme as earlier (**Figure 3e**), we see that the intermediately methylated reads tend to be on the hypermethylated allele (the presumed Xi) for bins with less than 40% DNAme and are on the hypomethylated allele for bins with greater than 50% DNAme. We further used this allelic DNAme to call XCI status, using thresholds at 25% and 75% DNAme per allele, with genes having both alleles below 25% being called as escaping from XCI and those having one allele below 25% and one above 75% being called as subject to XCI. These calls for SNPs within CpG islands had good agreement with previous calls with all 28 of the loci called escaping and 50/51 of the loci called subject to XCI being concordant. For SNPs outside of CpG islands the agreement was lower, with only 91/182 loci found escaping XCI and 235/274 loci found subject to XCI being concordant with their meta-status.

To explain the prevalence of intermediately methylated reads, we examined the DNAme per CpG across some of these islands where we observed that the DNAme level was not consistent (**Figure S6** for browser tracks across islands**, S7** for DNAme differences between adjacent CpGs). The intermediately methylated WGBS reads and CpG island DNAme averages seen above are likely due to this inter-CpG variability. We can also see this variability in DNAme in males although it is rarer. This problem seems exacerbated in the cancer samples examined here as compared to healthy samples from CREST. Examining the average DNAme difference between adjacent CpG sites we saw an average difference of 24% in cancer and 13% in healthy samples.

### A combined epigenetic model can predict XCI status across samples

DNAme has been shown repeatedly to be a strong predictor of XCI status, so we wanted to test whether the other epigenetic marks examined could also be used to predict XCI. Using simple thresholds to separate genes that have low values for a mark vs those with high values gave low accuracy (**Table S9**) with the most accurate mark being H3K27me3 with an accuracy of 68%, excluding genes called as variably escaping XCI. These solo histone thresholds particularly overcalled genes as variably escaping XCI, with a call as high as 87% of genes using gene-body H3K36me3. We moved to a random forest predictor model to predict the XCI status of genes using each individual mark per female sample along with matched male data. We trained on 75% of genes escaping XCI, using the remainder to test accuracy, and used twice as many genes subject to XCI for training. Using this predictor we could predict escape from XCI with accuracies ranging from 42% with H3K9me3 to 69% with H3K4me3 and for genes subject to XCI with accuracies ranging from 85% with gene-body H3K36me3 to 99% with H3K27ac (**Table S10**). In contrast, a similar model using CpG island DNAme data obtained a much better accuracy of 87% for predicting genes as escaping XCI and 99% for predicting genes as subject to XCI, showing the higher predictive ability of DNAme.

To get XCI status calls from histone mark data with an improved accuracy, we combined data from all of the histone marks and DNAme data from CEMT and trained a new random forest model [27]. This combined epigenetic XCI predictor was trained using XCI meta-status and was able to accurately predict genes escaping vs subject to XCI, with a median accuracy for genes outside the training set of 75% for genes escaping from XCI and 90% for genes subject to XCI (**Figure 4a**). We trained the model 20 separate times per sample and were confident in a prediction if 75%+ of the models agreed. A separate epigenetic XCI predictor was trained and used within each sample, however the models are capable of being used across samples within the same tissue with reduced accuracy and even across tissues (**Figure S8** for a summary of accuracies**, Figure S9** for comparisons between each tissue). Models in some tissues tended to overcall genes as subject to XCI while others overcalled genes as escaping from XCI. Across all samples, the model called 46 genes as escaping XCI, 780 genes as subject to XCI and seven genes as variably escaping from XCI (**Figure 4b, Table S4** for XCI status calls). While none of the genes predicted to escape XCI here have a meta-status of subject to XCI, 11 of the genes predicted to be subject to XCI have a meta-status of escaping XCI and an additional six genes are located in the PAR1 and are expected to escape XCI [2]. Comparing these predictions to our Xi/Xa expression based XCI status calls, 23 genes escape XCI in both sets while only two were called as escaping XCI by Xi/Xa expression and subject to XCI using this model and three genes had the opposite calls. Of the eight genes found variably escaping by Xi/Xa expression, three of them (CXorf38, PRKX and SMC1A) had their predicted XCI status across samples perfectly match that found by Xi/Xa expression. There are no genes that were given opposite calls across samples by our DNAme based calls and this model, however some of the genes found to variably escape differed between the two.

**Figure 4:**
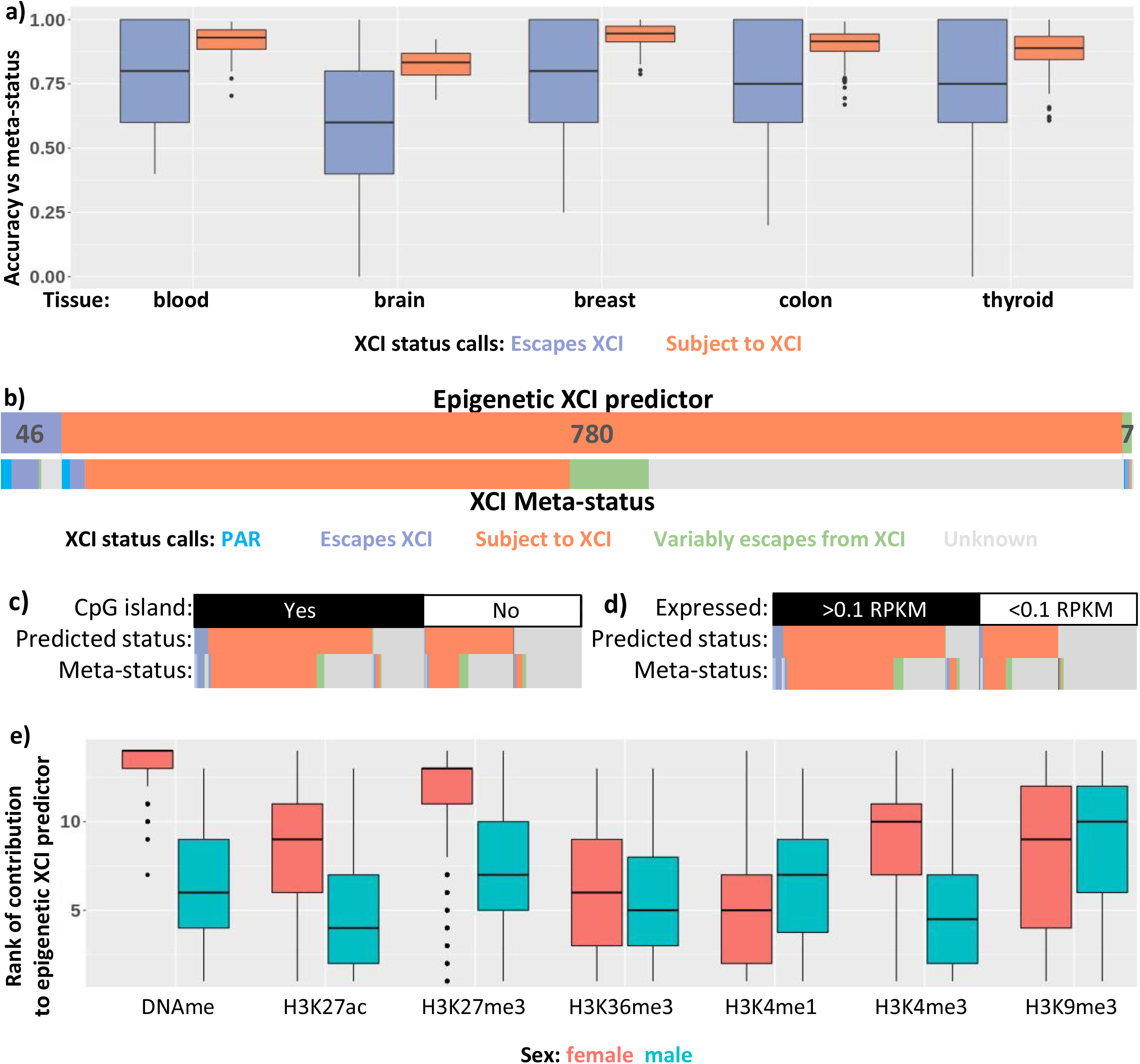
XCI status predictions with an epigenetic model expands the number of genes examineable. (a) Accuracy of our epigenetic predictor using DNAme and all six histone marks. Each point is one of the 20 models per sample. This accuracy is tested on genes outside of the training set. (b) The number of genes with each XCI status as predicted by our model, with their distribution by meta-status underneath. (c-d) As b, but further split by the presence of a CpG island (c) or by an expression threshold of 0.1 RPKM (d). (e) The predictive ability of each mark. Each mark was ranked per model on how important it was to the model, with the most important mark being ranked 14 and the least important being ranked 1. We used the marks within each female sample paired with the mean mark in similar male samples for the predictor, so both the female and male marks are featured here.

This epigenetic XCI predictor can predict XCI status across all genes on the X chromosome, without being limited by CpG density and levels of male DNAme that restricted the simple DNAme model; however, transcripts without high promoter CpG density are almost twice as likely to have inconsistent XCI status calls within the same sample while genes that are lowly expressed (median RPKM across samples <0.1) are over three times more likely to have an inconsistent XCI status call (**Figure 4c, d, Table S11** for the number of XCI status calls with and without expression and CpG islands). We predicted an XCI status for over 300 genes that did not have previously annotated XCI statuses, however ~200 of these had low expression and so may actually be silent on both the active and inactive X chromosome making an XCI status call moot. DNAme was the most important input for the models, with H3K27me3 being the next most important (**Figure 4d**).

In addition to the seven genes that our epigenetic predictor called as variably escaping XCI across samples, we predicted 48 genes having tissue-specific escape from XCI, and one gene with separate TSSs with opposite XCI status. We compared our epigenetic marks across samples, tissues and TSSs with opposite XCI statuses (**Table S12**). At genes predicted to variably escape across samples we found that very few marks had significant (t-test, adjusted p-value <0.01) differences between samples found escaping and those subject to XCI. DNAme was the exception to this with four of seven genes having significant DNAme differences. For the genes found variably escaping across tissues, all of the marks had multiple genes significantly different between tissues subject to XCI vs tissues escaping from XCI, but many of the genes that didn’t variably escape also had significant differences across tissues. Tissue-specific variable escape genes had significant enrichment (chi-square test, adjusted p-value<0.01) for genes with tissue-specific H3K27me3, H3K4me3, DNAme and expression over genes that did not variably escape from XCI. There was only one gene found to variably escape between TSSs so no statistical tests were possible, however there were differences between TSSs for H3K27ac, H3K4me1 and DNAme and at gene-body H3K9me3 for the different exons used.

Our initial thresholds to call variable escape across samples were arbitrary, so we decided to lower the percentage of samples with each XCI status required to call a gene as variably escaping from XCI. At our threshold requiring 33% of samples to have each XCI status in order to be called as variably escaping from XCI, we found 7 of 1155 genes to be variably escaping. Lowering this threshold to 25% found 35 variably escaping genes, at 10% we found 304 genes and at 5% we found 476 genes. This shows that there is no natural threshold at which genes become variable, rather a large proportion of genes will occasionally differ in their XCI status, but few genes are highly variable across samples. As the threshold for calling genes as variably escaping decreased, the percentage of these genes with significant DNAme differences between samples with opposite XCI statuses decreased down to 20% and the percentage of genes with H3K27me3 differences rose to 27% (**Table S13**); however, we must also consider that the cancer origin of these samples may contribute to rare epigenetic mis regulation.

To validate our conclusions from this model on healthy samples, we trained our overall epigenetic predictor on the CREST dataset. The CREST dataset contains nine samples for which we were able to obtain all the required epigenetic data for our predictor. We predicted 88 genes escaping from XCI, 802 subject to XCI, 40 variably escaping across samples, ten across tissues and six across TSSs. These calls are similar to those in the CEMT data, with 95% of genes with calls from both datasets agreeing **(Table S14**). The genes variably escaping from XCI in the CEMT dataset tended to be escaping XCI in CREST while genes variably escaping in CREST tended to be subject to XCI in the CEMT dataset. The number of genes variably escaping from XCI is increased in CREST, possibly due to how few samples were required for variable escape (three with each XCI status) decreasing stringency. Another possibility is that having random Xi choice doubles the chance of seeing variability in XCI status and the predictor may be sensitive to a change in XCI status in only half of the cells. The number of tissue-specific genes is much reduced in CREST however, likely due to having only two tissues rather than five. Very few of the genes variably escaping across individuals in CREST had significant differences between samples subject to XCI and those escaping from XCI (**Table S15**). CREST tissue-specific genes had significant differences in H3K27me3, DNAme and expression between tissues, all three of which were also significant in CEMT samples. CREST had enough genes variably escape across TSSs to see that H3K4me3, H3K27me3 and DNAme were significantly different between TSSs escaping and TSSs subject to XCI in females. Males had significant differences in H3K4me3, H3K27ac, H3K27me3, H3K36me3 and DNAme between TSSs escaping vs subject to XCI in females, which suggests that these TSS also differ significantly on the Xa. These TSSs may be predisposed to have different XCI statuses based on their epigenetic landscape prior to XCI or the Xa differences may be misleading the predictor causing it to predict different XCI statuses.

To test whether the variably escaping genes found by our CEMT predictor intrinsically differ from genes with a consistent XCI status, we trained a pair of models, the first to differentiate genes that are constitutively escaping XCI across all samples from genes that are escaping XCI in the sample but were found to variably escape from XCI across all samples. The second model was trained to differentiate genes which were consistently predicted to be subject to XCI from those which were predicted as being subject to XCI within the sample but variably escape across all samples. To have enough variably escaping genes for these models, we used the 10% variable escape threshold. The model differentiating constitutive escape from variable escape genes overcalled variable escape (median accuracy of 75%) but all other accuracy metrics were over 85%, while the model differentiating genes subject to XCI from genes variably escaping from XCI performed worse, overcalling genes as subject to XCI with a median accuracy across samples of only 15% for genes called as variably escaping from XCI (**Figure S10**). This could suggest that many of these variably escaping genes are just genes subject to XCI that have been miscalled, or it could be that they are epigenetically closer to genes subject to XCI. These genes were depleted for meta-status calls of subject to XCI compared to the overall distribution of meta-statuses; however, as 70% of genes had a meta-status call of being subject to XCI, there were still more predicted variably escaping genes with a meta-status call of subject to XCI than any other XCI status. These variably escaping genes were predicted to be subject to XCI in more samples than they were predicted to escape from XCI (median of 18 samples subject to XCI and seven escaping from XCI) (**Figure S11**). The models differentiating genes escaping XCI from those variably escaping XCI tended to rely on DNAme, H3K27ac and H3K4me3, in both the female samples and their matched male controls while those differentiating genes subject to XCI from those variably escaping XCI tended to rely equally on all marks except H3K36me3 and H3K9me3 (**Figure S12**).

### Independent regulation of variable escape across a region

As an application of our epigenetic XCI predictor, we examined XCI status calls per sample across a region that is enriched in genes variably escaping from XCI according to their meta-status (**Figure 5a**) [2]. We did this to understand the scale at which variably escaping genes are regulated. We found that many of the genes in this region that are annotated as variably escaping from XCI had low levels of variable escape with few samples differing from the most common XCI status. The genes that vary in XCI status across samples change their XCI status independent of the XCI status of neighboring genes, suggesting that regulation of variably escaping genes happens at the single gene level and not at the domain level. Additionally, we saw genes that had multiple TSSs with different XCI statuses and genes that are bidirectional from the same promoter with opposite XCI status showing that the scale of regulation could be narrowed even further. All of the genes in this region that showed variable escape here, except for IRAK1, had significant differences for some combination of marks including H3K9me3, H3K27me3 and DNAme between genes escaping vs subject to XCI (p-val<0.05, **Figure 5b, S13** for which marks were significant per TSS). Euchromatic marks were less frequently seen to be significantly different.

**Figure 5:**
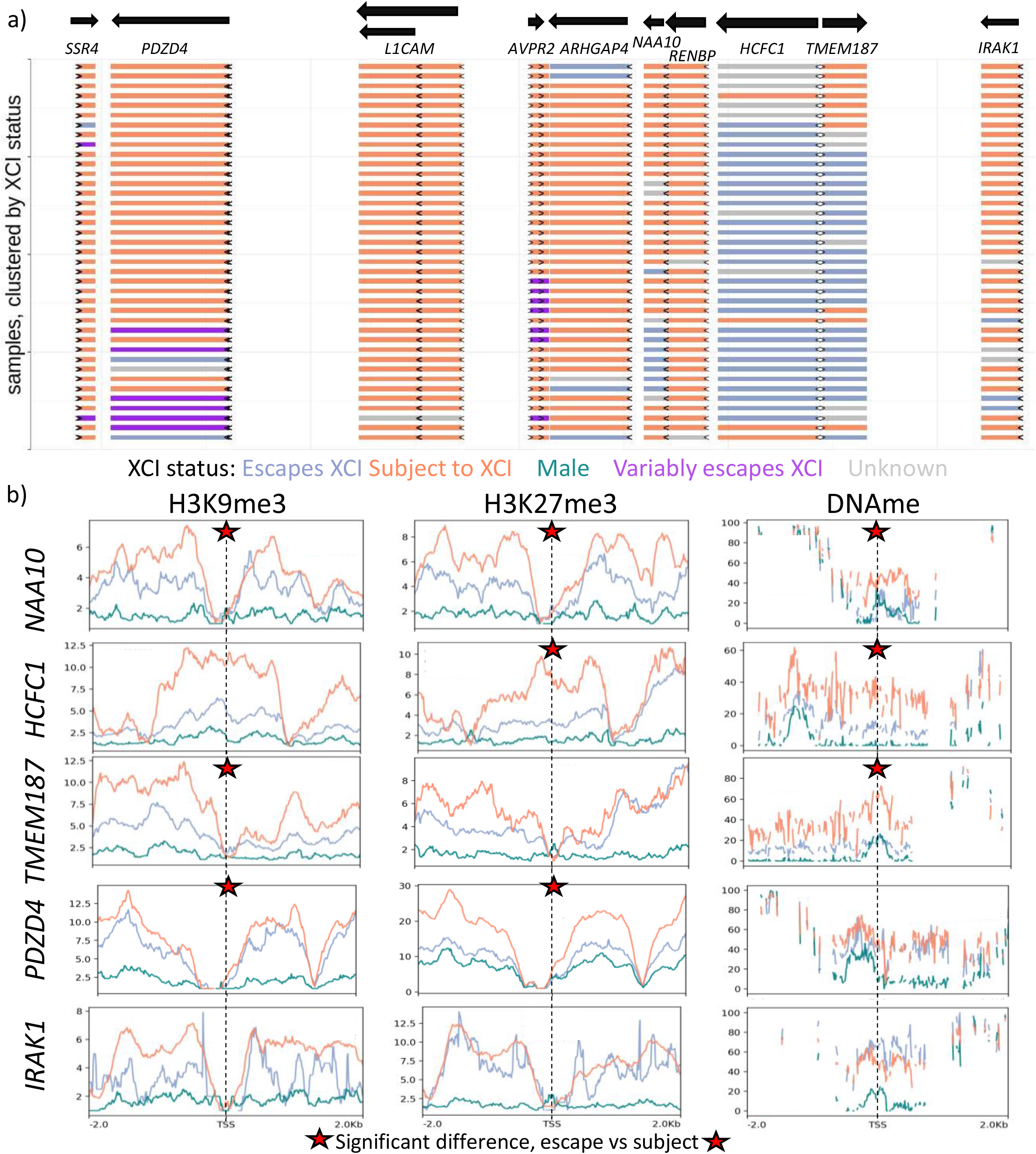
XCI status calls are independent between neighboring variably escaping genes. (a) A map of a variably escaping region, with genes colored by their XCI status as predicted by our random forest model using all epigenetic marks available. The samples were clustered based on their XCI status calls within the region. Arrows indicate where each TSS is located, and they point in the direction of transcription. (b) Metagene plots for the epigenetic marks that were most commonly significantly different between samples subject to XCI vs those escaping from XCI at the above variably escaping genes. Genes were chosen to show every combination of which mark is significant per gene, that we saw in this region. Marks that were significant at a gene are marked with a star.

## Discussion

XCI is a classic paradigm for epigenetic regulation, yet why some genes are resistant to silencing (or the maintenance of silencing) and escape XCI remains unresolved. Here we have examined the epigenetic differences between genes escaping and those subject to XCI. Epigenetic marks tended to be more different between males and females than between genes escaping vs subject to XCI. Genes escaping XCI tended to have similar epigenetic marks between the Xi and Xa, except for H3K27me3 which is higher on the Xi. The increased Xi H3K27me3 at genes escaping XCI may be why escape genes can have as low as 10% expression from the Xi compared to the Xa; while the Xa-like status at other marks would allow some expression to continue. Genes subject to XCI tend to have higher heterochromatic marks on the Xi and lower euchromatic marks, which supports these genes not being expressed on the Xi. Enhancers were also seen to show strong male:female differences that mostly reflected the presence of the Xi with a lesser contribution from inactivation status. Other studies have seen a general enrichment of heterochromatic marks at genes subject to XCI and euchromatic marks enriched at genes escaping from XCI (reviewed in [13]).

Across all our epigenetic analyses, DNAme stood out as being the most reflective of a gene’s XCI status. The euchromatic mark H3K4me3 was the histone mark that was most significant for differentiating genes escaping from those subject to XCI, while the heterochromatic mark H3K27me3 had the largest Xi:Xa difference and was the most predictive histone mark for our epigenetic predictor. A previous study, which used a random forest model to predict XCI statuses and silencing timing in mice, found that DNAme often ranked below many of their histone marks, including H3K27ac, H3K4me1 and H3K27me3 [23]. In addition to the possible species differences, their model may not rely on DNAme as much due to them including numerous genomic features and transcription factor binding annotations, with distance to *Xist* and gene density being their top two features. We chose not to incorporate more features as we wanted to find XCI differences across samples, and therefore wanted to only include features that were sample-specific. Our model however does not account for the interaction genomic features may have with the epigenetic marks examined here. If we had both genetic and epigenetic data for a large number of samples, that would have been ideal for determining the genetic and epigenetic determinants of XCI.

In this study we used multiple different methods to predict the XCI status of genes and examined how different epigenetic marks changed across genes with differing XCI statuses. We found similar distributions of genes escaping, variably escaping or subject to XCI across our DNAme analyses as our previous Xi/Xa expression analysis [12], while our epigenetic predictor predicted twice as many genes as subject to XCI with similar levels of genes escaping and variably escaping from XCI. A large proportion of the additional genes found subject to XCI by our epigenetic predictor may in fact be silenced on both the Xa and Xi, as 68% of them had a median expression across samples under 0.1 RPKM.

The threshold at which to call genes as ‘variable’ in XCI status is arbitrary. We used a threshold requiring 33% of samples to have each XCI status to call variable escape from XCI in our DNAme and epigenetic predictors as used previously [9, 12], with the greater number of samples with each XCI status improving the power of our statistical tests comparing epigenetic marks across samples with opposite XCI statuses. Decreasing the threshold increased the number of genes variably escaping from XCI and the number of epigenetic marks that were significant in at least one gene, but decreased the percentage of genes significant for DNAme that was the only mark ever significant for over 50% of genes in a dataset.

We observed that variable escape from XCI was regulated at the level of single genes, with adjacent genes varying their XCI status independently. In contrast, a study in mice found clusters of genes that variably escape across their three cell lines, with adjacent genes often having the same XCI status across lines [10]. They also found that these clusters colocalize with TADs, with one line having the majority of a TAD escaping XCI and another line having only part of it escaping. An interesting candidate regulator of regional control is *SMCHD1*. In mice with *SMCHD1* knocked-out, regions enriched with variably escaping genes were upregulated, while genes that constitutively escaped from XCI were not affected; however, no impact was seen on variable escape genes in human patients with heterozygous *SMCHD1* mutations [28]. Another study found variants with low expression of *SMCHD1*, *ZSCAN9* and *HBG2/TRIM*6 associated with hypomethylation of X-linked CpG islands, with affected islands enriched near genes that variably escape from XCI [29]. Overall, there is evidence for both domain-level and gene-specific regulation of escape. We suggest that for some domains the former predominates, while for other genes the latter predominates. Additionally, the domain featured in Figure 4 (and other variably escaping domains) are at a threshold where individual genes within the domain can have either XCI status based on local factors.

One drawback to this study is that many of our results relied on cancer datasets that may have differences from healthy tissues and epigenetic instability. DNA methyltransferases and histone modifying enzymes are commonly mutated in cancer, and 5-10% of CpG islands that should be unmethylated become methylated (reviewed in [30]). We would expect the changes from epigenetic instability to differ between cancers and cause more genes to variably escape from XCI, however we saw a similar number of genes variably escaping from XCI in the CEMT cancer dataset as in the healthy CREST dataset. Despite these problems, we used the CEMT dataset because it had a standardized set of epigenetic marks across many samples and the clonality of cancer allowed us to examine expression and DNAme allelically. We found that other datasets, did not always have all the marks from the same samples, were lacking females or sex labels or had mislabeled sex.

We had some discordant calls between the various methods employed here. Genes could be falsely called as subject to XCI in the Xi/Xa expression-based analysis if the alternate SNP allele no longer mapped to the same region or if heterozygosity was miscalled. DNAme has been seen mis regulated in many cancers [30]. The cancer cells could have mutations mosaic between the parts sampled for different analyses. Our epigenetic predictor did not obtain 100% accuracy on its training data so we expect some of the calls made with it to be false, while the training data could also have false calls further hurting its ability to make accurate XCI status calls.

## Conclusions

Our study has shown that most of the epigenetic marks assayed (H3K4me1, H3K4me3, H3K9me3, H3K27ac, H3K27me3, H3K36me3, and DNAme) had male-female differences on the X chromosome, while fewer marks were significantly different at genes with opposite XCI statuses within females. To account for dosage differences, we calculated the contributions of the Xa and Xi to these sex-biased modifications. Genes subject to XCI had higher heterochromatic marks on the Xi and lower euchromatic marks on the Xi while genes escaping XCI tended to have equal levels of marks on the Xa and Xi, except for H3K27me3 that was high on the Xi. Genes that escape from XCI are not expressed at 100% of the level of the Xa, which supports this conclusion. No mark other than DNAme was very accurate at predicting XCI status; however, combining all of the epigenetic marks together allowed us to call XCI status for genes without CpG islands, where DNAme alone is unable to establish a call. Most marks were significantly different between samples escaping vs subject to XCI at variably escaping genes, but which marks were significant was not consistent between genes and no mark was significant across all the genes. This may be due to variably escaping genes having multiple ways in which they are regulated. DNAme intermediate to what is expected for genes escaping vs subject to XCI is enriched at variably escaping genes and is mostly due to inconsistent DNAme on the Xi. Neighboring variably escaping genes were seen to regulate their XCI status independently from each other, suggesting local regulatory elements. Overall, we see that escape from XCI is influenced by chromatin modifications that can be independent of each other. Understanding how genes escape from XCI will further our understanding of epigenetics in general, and may allow us to control which genes are escaping from XCI and rescue X-linked mutations in females.

## Methods

### Previous XCI status calls

We used XCI meta-status calls from [2] for all comparisons with past XCI statuses and to train our models. Genes that escape and mostly escaped were combined together due to the small size of these categories, with genes in the PAR1 being left out or having their own separate category depending on the analysis. Genes that were mostly subject to XCI were combined with genes subject to XCI for comparisons between studies but were left out when training models. Genes that were annotated as variably escaping, mostly variably escaping and discordant across studies were combined together as variably escaping genes for comparisons here.

### Histone ChIP-seq analysis

Histone ChIP-seq bigwig files were downloaded from the IHEC data portal [31] and quantified with bigWigAverageOverBed [32] for a region 500bp upstream of TSSs as annotated by Gencode [33]. We normalized the data across samples by multiplying samples to have the same total depth (including all chromosomes). The metagene plots for figure 5 and S2 were generated using Deeptools computeMatrix and plotProfile [34].

### Expression analysis

Xi/Xa expression based XCI status calls per sample were generated previously [12]. In this study we used a different threshold to identify variably escaping genes, requiring at least two samples with each XCI status. This narrows the number of variably escaping genes and increases the chance that those found would have enough samples to reach significance. The overall expression level of genes was calculated using bigwig files downloaded from the CEEHRC data portal [35] and quantified as RPKM using VisRseq [36].

### DNAme analysis

WGBS bigwig files were downloaded from the IHEC data portal [31] and quantified with bigWigAverageOverBed [32] for a region 500bp upstream of TSSs as annotated by Gencode [33]. DNAme thresholds established in [12] were used to determine which genes were escaping XCI and which were subject to XCI. These thresholds are: DNAme<10% escapes XCI, 15%<DNAme<60% subject to XCI, and DNAme>60% hypermethylated. A threshold of DNAme<15% in males was used to filter out TSSs that were methylated on the Xa and therefore not informative for this analysis. To see the differences between adjacent CpGs, we converted bigWig files to bedGraphs and for each island we used R to find the mean absolute value difference between each adjacent CpG.

DNAme per read was calculated by downloading WGBS bam files and using a script to count the number of unmethylated and methylated CG dinucleotides per read within CpG islands within 2kb of TSSs. For allelic DNAme, we did similar but only examined reads that overlapped heterozygous SNPs identified in our Xi/Xa analysis and had to reconstruct the read from the CIGAR string in the bam file in order to determine the allele of origin. We analyzed allelic DNAme for SNPs within 2kb of TSSs, and noted which were found in CpG islands.

We used bins for every 10% increase in mean DNAme, and chose bins for each individual gene per sample separately. All of the reads per bin, across all genes and female samples were used for figure 3c. For allelic DNAme we first filtered out polymorphisms where the alleles were CT or GA as bisulfite conversion makes it impossible to differentiate these. We then combined all reads with a C or T allele and all reads with a G or A allele together and filtered out polymorphisms without at least five of each allele type in a sample. The mean DNAme per read per allele type was then calculated and this was used to make XCI status calls per polymorphism in each sample with enough reads. The thresholds were 0.25 and 0.75 for our XCI status calls with polymorphisms with both alleles below 0.25 being called as escaping from XCI and both alleles higher than 0.75 being called as hypermethylated. Polymorphisms with one allele above 0.75 and the other allele below 0.25 were called as subject to XCI. The DNAme per read per polymorphism was binned as above, but instead of using the mean DNAme across all reads, we determined the mean DNAme per allele and used the mean of that; this was done so that we get the mean between the Xi and Xa if there are more reads for one than the other. Additionally, we determined which allele was lower for each polymorphism and graphed the low allele separately from the high allele, per bin.

### Histone-based XCI status predictions

A simple histone predictor was made using genes with known XCI status as published in [2], and defining XCI status for genes within two standard deviations of the mean for each XCI status, similar to a model used in [20]. Because the mean of genes subject to XCI and the mean of genes escaping XCI were often within two standard deviations of each, the average of these two means was often used as a threshold instead.

For our random forest models, we wanted to include both male and female data, and breast did not have any male data so we used the kmeans function in R to cluster all of our samples based on autosomal levels of all seven epigenetic marks used herein. With three clusters we had multiple male and female samples in each cluster. As input for our models, we used individual female data per sample and matched it with the mean values per gene across males in the same cluster.

Random forest models were trained using the R package caret [27] with the trainControl method cv and the train method rf. We trained the model on genes known to escape or be subject to XCI [2]. The training metric was ROC, tunelength was 5 and ntree was 1500. Three genes escaping and subject to XCI were left out of the training set and used to check accuracy of overall calls. We trained twenty models per sample, with each model being trained on a random sample of 75% of the genes escaping XCI and twice as many genes subject to XCI, with each iteration of the model using 75% of the number of input escaping genes. Accuracy per model was tested on the remaining genes with known XCI status. Genes were considered as escaping or subject to XCI if 15+ of 20 models predicted them as escaping or subject to XCI respectively. Separate categories were made for genes where only 12-14 of the models agreed on the gene’s XCI status, being annotated as leaning subject or leaning escape. Overall calls were made across samples with genes with 66% or more of samples agreeing on a gene’s XCI status being called as subject to or escaping from XCI, genes with at least 33% or more of all samples having each XCI status being called as variably escaping from XCI, and genes that required the leaning categories to reach 66% of samples having a status being annotated with a similar leaning status.

### Statistical comparisons

All statistical comparisons were done in R [37]. The majority were t-tests with a Benjamini-Hochberg (BH) multiple testing correction [38] with results deemed significant if they had an adjusted p-value<0.01. The one test with a different threshold was for comparing genes variably escaping XCI as determined by Xi/Xa expression. This test used a 0.05 threshold and had no multi testing correction due to a low sample size, with most genes only having 2 or 3 of each category. If we had a larger sample size, a more stringent test would be preferred. We also used a chi-square test to determine enrichment of significant histone differences between tissues and TSSs, with p-value of 0.01.

## Supporting information

Supplemental Figures and small tables

## List of abbreviations

BH: Benjamini-Hochberg
CEMT: Center for Epigenome Mapping Technologies
ChIP-seq: chromatin immunoprecipitation sequencing
CREST: Core Research for Evolutional Science and Technology
DNAme: DNA methylation
IHEC: International Human Epigenome Consortium
meta-status: XCI status calls from Balaton et al., 2015
PAR: pseudo-autosomal region
RNA-seq: RNA sequencing
TAD: topologically associating domain
TSS: transcription start site
WGBS: whole genome bisulfite sequencing
Xa: active X
XCI: X-chromosome inactivation
Xi: inactive X

## Acknowledgements

We thank the other members of the Brown lab for helpful comments during the development of this project.

Most of the analyses conducted here used data generated by The Canadian Epigenetics, Epigenomics, Environment and Health Research Consortium (CEEHRC) initiative funded by the Canadian Institutes of Health Research (CIHR), Genome BC, and Genome Quebec. Information about CEEHRC and the participating investigators and institutions can be found at http://www.cihr-irsc.gc.ca/e/43734.html. We would also like to thank the research groups which generated the other sources of data used in this analysis.

## Table Legends

**Table S1: List of samples used.** See additional file 1. For CEMT samples, tissue was manually annotated to combine samples from related areas. Columns D through L are true if the dataset was available for the sample. Patient health status and sample disease are the annotations done by CEMT. CREST samples were only used for the epigenetic predictor and only samples with all datasets available were included here.

**Table S2: Comparison of histone marks between sex and XCI status.** See additional file 2. The first two sheets show BH adjusted p-values comparing female vs male and escape genes vs those subject to XCI per mark in in CEMT with our meta-status and Xi/Xa expression based XCI status calls, along with CREST data with meta-status calls and CREST data at enhancers with meta-status calls of linked genes. The last 2 sheets show the median value per mark with each sex and XCI status on the left and on the right shows the Xi/Xa ratio and log2 fold change per mark calculated based off of that median.

**Table S3: The ratio of TSSs with significant differences between males and females for various epigenetic marks using CEMT data.** The denominator was the total number of informative TSSs for which we had data. For most marks this was measured as 500bp upstream of the promoter, but for H3K36me3 we measured the mark across exons. For H3K36me3 we used unique transcripts instead of unique TSSs. Marks significant in over 70% of informative TSSs are in bold. All of the H3K27me3 data from ENCODE was downloaded and used as a replication dataset. Chromosome 7 (chr7) was included as an example autosome.

**Table S4: All XCI status calls made here.** See additional file 3. The first sheet contains a single XCI status call per gene per method. Published calls are from Balaton, et al. 2015. Other sheets contain all calls per sample for each method. Each row is one entry into the model, so Xi/Xa is per gene and the others are for unique transcripts. For DNAme, the samples on the far right in shades of grey are males while the samples on the left in color are females. For the epigenetic predictor, separate low confidence categories were made for when transcripts have only 12-14 of the 20 models per sample predicted a certain XCI status.

**Table S5: Significance of differences in epigenetic marks between samples with opposite XCI statuses at genes found variably escaping XCI by Xi/Xa expression.** We also tested whether expression differed between samples with opposite XCI status. Presented here are the p-values of t-tests. Those with p-values less than 0.05 are in bold. nE and nS are the number of samples escaping or subject to XCI for each gene.

**Table S6: The differences in epigenetic marks between samples with opposite XCI statuses at genes found variably escaping XCI by Xi/Xa expression.** The mean value for samples subject to XCI was subtracted from the mean value for samples escaping XCI. Those found significant in Table 1 are bolded. Genes with multiple transcripts are included multiple times, even if they share a TSS.

**Table S7: adjusted p-values comparing marks in females between genes found subject to XCI vs escaping XCI by DNAme.** Those in bold are significant (adjusted p-value<0.01).

**Table S8: Distribution summary for DNAme per read.** The number is what percent of reads in each bin were below 25%, between 33 and 66% or over 75% DNAme.

**Table S9: The accuracy of simple models predicting XCI status from a single histone mark.** These accuracies are low because the models overpredicted variable escape from XCI as there is large overlap between the two XCI statuses.

**Table S10: The accuracy of random forest models predicting XCI status from a single histone mark**.

**Table S11: XCI status calls made using a random forest epigenetic predictor, split by presence or absence of a CpG island and expression.** The threshold used to split low from high expression is a median of 0.1 RPKM across samples. Inconsistent predictions had over a third of samples with fewer than 15 of the 20 models trained agree on an XCI status.

**Table S12: The percent of genes found variably escaping by our epigenetic predictor with significant differences in various epigenetic marks.** Genes were counted as significant if BH corrected p-values were less than 0.01 when comparing samples predicted as subject to XCI to samples predicted as escaping from XCI. The total number of genes row shows the total number of genes in each category. The variable escape across tissues and TSSs categories have 2 columns each, the left column being the percent of variably escaping genes with significant differences between tissues/TSSs and the right column being the percent of all genes on the X chromosome with differences between tissues/TSSs. Highlighted in blue are marks that were significantly more likely to have significant differences between tissues/TSSs at genes predicted to variably escape than in all X linked genes.

**Table S13: The percent of genes found variably escaping by our epigenetic predictor with significant differences in various epigenetic marks across various variable escape thresholds.** Variable escape threshold is the number of samples with each XCI status (escaping from XCI and subject to XCI) that were required in order to call a gene as variably escaping from XCI across samples. Genes were counted as significant if BH corrected p-values were less than 0.01 when comparing samples predicted as subject to XCI to samples predicted as escaping from XCI.

**Table S14: Comparing XCI status calls made by an epigenetic predictor in the CEMT dataset vs a similar model in the CREST dataset**.

**Table S15: The percent of genes found variably escaping by our epigenetic predictor in the CREST dataset with significant differences in various epigenetic marks.** Genes were counted as significant if BH corrected p-values were less than 0.01 when comparing samples predicted as subject to XCI to samples predicted as escaping from XCI. The total number of genes row shows the total number of genes in each category. The variable escape across tissues and TSSs categories have 2 columns each, the left column being the percent of variably escaping genes with significant differences between tissues/TSSs and the right column being the percent of all genes on the X chromosome with differences between tissues/TSSs. Highlighted in blue are marks that were significantly more likely to have significant differences between tissues/TSSs at genes predicted to variably escape than in all X linked genes.

---

**Figure S1: log2(Xi/Xa) for epigenetic marks in CREST at promoters.** Data from CREST is shown. Significance for the various t-tests featured in Table S2 are shown by the differently colored star.

**Figure S2: Meta-gene plots of histone marks within 50kb of genes, separated by XCI status**. The plots were generated using deeptools computeMatrix and plotProfile on bigwig files that were the mean across samples. Lighter shaded regions show the standard deviation of each mark.

**Figure S3: log2(Xi/Xa) for epigenetic marks in CEMT at enhancers mapping to genes that escape from or are subject to XCI.** Enhancers are split by whether they are located within a gene (genic) or not (intergenic).

**Figure S4: Expression across exons for genes with significantly different expression in samples with opposite XCI statuses.** XCI status per sample was determine here using Xi/Xa expression.

**Figure S5: Differences in epigenetic marks between samples found escaping vs subject to XCI at variably escaping genes in DNAme.** For most of these marks, the region 500bp upstream of the promoter is used, except for H3K36me3 which uses the gene body.

**Figure S6: IGV view of DNAme bigwig tracks at two variably escaping genes.** a) A view of the CpG island at CITED1. b) a view of the CpG island at NAA10. A broad representation of samples was sought, some hypomethylated, some hypermethylated and some inconsistent across the CpG island. Broad hypermethylation in males at these genes was rare but is included here as an example of an extreme.

**Figure S7: average DNAme difference between adjacent CpGs per CpG island.** Each point is the average DNAme difference between adjacent CpGs for an individual island, averaged again across samples. Islands are colored by the meta-status of the closest TSS within 2kb. Chr7 was chosen as an autosomal control to show whether the differences are X specific. Males and females from CEMT were used to check for sex specificity and females from CREST were included to check for cancer specificity.

**Figure S8: Accuracy when models trained in one sample are tested on other models**.

**Figure S9: Accuracy when models trained in one sample are tested on other models, separated per tissue comparison.** The numbers at the bottom of each plot are the median accuracy. Each point is the accuracy at predicted an XCI status when a model from the training tissue on a sample in the predicted tissue. Eaccuracy is accuracy at predicting genes as escape from XCI. Saccuracy is accuracy when predicting genes as subject to XCI.

**Figure S10: Accuracy metrics when predicting which genes variably escape from XCI across samples using data from individual samples.** On the left are metric when a model is trained on only genes called as escaping XCI in that sample, while the right is metrics when a model is trained on only genes called as subject to XCI. VE is variably escaping from XCI. Subject is subject to XCI.

**Figure S11: The number of samples called as escaping vs subject to XCI per transcript by our epigenetic predictor**.

**Figure S12: Ranked importance of the marks used to predict which genes variably escape across samples.** The contributions to each model from each mark were ranked, with rank 14 being the most important and rank one being the least important.

**Figure S13: Which marks were significantly different between samples predicted as escaping vs subject to XCI in a variably escaping region.** Transcript ID is the order that the transcripts are located along the chromosome. There are multiple transcripts per gene but they may be sharing the same TSS and have the same data for all marks but H3K36me3. Vertical lines are drawn denoting which transcripts belong with each gene.

## References

1. Balaton BP, Dixon-McDougall T, Peeters SB, Brown CJ. The eXceptional nature of the X chromosome. Hum Mol Genet. 2018;27:R242–9.

2. Balaton BP, Cotton AM, Brown CJ. Derivation of consensus inactivation status for X-linked genes from genome-wide studies. Biol Sex Differ. 2015;6:35.

3. Carrel L, Willard HF. X-inactivation profile reveals extensive variability in X-linked gene expression in females. Nature. 2005;434:400–4.

4. Dunford A, Weinstock DM, Savova V, Schumacher SE, Cleary JP, Yoda A, et al. Tumor-suppressor genes that escape from X-inactivation contribute to cancer sex bias. Nat Genet. 2017;49:10–6.

5. Navarro-Cobos MJ, Balaton BP, Brown CJ. Genes that escape from X-chromosome inactivation: Potential contributors to Klinefelter syndrome. Am J Med Genet C Semin Med Genet. 2020;184:226–38.

6. Tukiainen T, Villani A-C, Yen A, Rivas MA, Marshall JL, Satija R, et al. Landscape of X chromosome inactivation across human tissues. Nature. 2017;550:244–8.

7. Godfrey AK, Naqvi S, Chmátal L, Chick JM, Mitchell RN, Gygi SP, et al. Quantitative analysis of Y-Chromosome gene expression across 36 human tissues. Genome Res. 2020;30:860–73.

8. Helena Mangs A, Morris BJ. The Human Pseudoautosomal Region (PAR): Origin, Function and Future. Curr Genomics. 2007;8:129–36.

9. Carrel L, Willard HF. X-inactivation profile reveals extensive variability in X-linked gene expression in females. Nature. 2005;434:400–4. doi:10.1038/nature03479.

10. Marks H, Kerstens HHD, Barakat TS, Splinter E, Dirks RAM, van Mierlo G, et al. Dynamics of gene silencing during X inactivation using allele-specific RNA-seq. Genome Biol. 2015;16:149.

11. Goto Y, Kimura H. Inactive X chromosome-specific histone H3 modifications and CpG hypomethylation flank a chromatin boundary between an X-inactivated and an escape gene. Nucleic Acids Res. 2009;37:7416–28.

12. Balaton BP, Fornes O, Wasserman WW, Brown CJ. Cross-species examination of X-chromosome inactivation highlights domains of escape from silencing. Genetics. 2020.

13. Balaton BP, Brown CJ. Escape Artists of the X Chromosome. Trends Genet. 2016;32:348–59.

14. Berletch JB, Ma W, Yang F, Shendure J, Noble WS, Disteche CM, et al. Escape from X inactivation varies in mouse tissues. PLoS Genet. 2015;11:e1005079.

15. Vacca M, Della Ragione F, Scalabrì F, D’Esposito M. X inactivation and reactivation in X-linked diseases. Semin Cell Dev Biol. 2016;56:78–87.

16. Mengel-From J, Lindahl-Jacobsen R, Nygaard M, Soerensen M, Ørstavik KH, Hertz JM, et al. Skewness of X-chromosome inactivation increases with age and varies across birth cohorts in elderly Danish women. Sci Rep. 2021;11:4326.

17. Larson NB, Fogarty ZC, Larson MC, Kalli KR, Lawrenson K, Gayther S, et al. An integrative approach to assess X-chromosome inactivation using allele-specific expression with applications to epithelial ovarian cancer. Genet Epidemiol. 2017;41:898–914.

18. Moreira de Mello JC, Fernandes GR, Vibranovski MD, Pereira LV. Early X chromosome inactivation during human preimplantation development revealed by single-cell RNA-sequencing. Sci Rep. 2017;7:10794.

19. Hagen SH, Henseling F, Hennesen J, Savel H, Delahaye S, Richert L, et al. Heterogeneous Escape from X Chromosome Inactivation Results in Sex Differences in Type I IFN Responses at the Single Human pDC Level. Cell Rep. 2020;33:108485.

20. Cotton AM, Price EM, Jones MJ, Balaton BP, Kobor MS, Brown CJ. Landscape of DNA methylation on the X chromosome reflects CpG density, functional chromatin state and X-chromosome inactivation. Hum Mol Genet. 2015;24:1528–39.

21. Qu K, Zaba LC, Giresi PG, Li R, Longmire M, Kim YH, et al. Individuality and variation of personal regulomes in primary human T cells. Cell Syst. 2015;1:51–61.

22. Kucera KS, Reddy TE, Pauli F, Gertz J, Logan JE, Myers RM, et al. Allele-specific distribution of RNA polymerase II on female X chromosomes. Hum Mol Genet. 2011;20:3964–73.

23. Barros de Andrade E Sousa L, Jonkers I, Syx L, Dunkel I, Chaumeil J, Picard C, et al. Kinetics of -induced gene silencing can be predicted from combinations of epigenetic and genomic features. Genome Res. 2019;29:1087–99.

24. Wang Z, Willard HF, Mukherjee S, Furey TS. Evidence of influence of genomic DNA sequence on human X chromosome inactivation. PLoS Comput Biol. 2006;2:e113.

25. Barski A, Cuddapah S, Cui K, Roh T-Y, Schones DE, Wang Z, et al. High-resolution profiling of histone methylations in the human genome. Cell. 2007;129:823–37.

26. Fishilevich S, Nudel R, Rappaport N, Hadar R, Plaschkes I, Iny Stein T, et al. GeneHancer: genome-wide integration of enhancers and target genes in GeneCards. Database. 2017;2017. doi:10.1093/database/bax028.

27. Website. Max Kuhn (2020). caret: Classification and Regression Training. R package version 6.0-86. https://CRAN.R-project.org/package=caret. Accessed 11 Dec 2020.

28. Wang C-Y, Brand H, Shaw ND, Talkowski ME, Lee JT. Role of the Chromosome Architectural Factor SMCHD1 in X-Chromosome Inactivation, Gene Regulation, and Disease in Humans. Genetics. 2019;213:685–703.

29. Luijk R, Wu H, Ward-Caviness CK, Hannon E, Carnero-Montoro E, Min JL, et al. Autosomal genetic variation is associated with DNA methylation in regions variably escaping X-chromosome inactivation. Nat Commun. 2018;9:3738.

30. Dawson MA, Kouzarides T. Cancer epigenetics: from mechanism to therapy. Cell. 2012;150:12–27.

31. Bujold D, Morais DA de L, Gauthier C, Côté C, Caron M, Kwan T, et al. The International Human Epigenome Consortium Data Portal. Cell Syst. 2016;3:496–9.e2.

32. Kent WJ, Zweig AS, Barber G, Hinrichs AS, Karolchik D. BigWig and BigBed: enabling browsing of large distributed datasets. Bioinformatics. 2010;26:2204–7.

33. Harrow J, Frankish A, Gonzalez JM, Tapanari E, Diekhans M, Kokocinski F, et al. GENCODE: the reference human genome annotation for The ENCODE Project. Genome Res. 2012;22:1760–74.

34. Ramírez F, Ryan DP, Grüning B, Bhardwaj V, Kilpert F, Richter AS, et al. deepTools2: a next generation web server for deep-sequencing data analysis. Nucleic Acids Res. 2016;44:W160–5.

35. Canadian Epigenomes. http://www.epigenomes.ca/data-release/hg38/. Accessed 14 Aug 2020.

36. Younesy H, Möller T, Lorincz MC, Karimi MM, Jones SJM. VisRseq: R-based visual framework for analysis of sequencing data. BMC Bioinformatics. 2015;16 Suppl 11:S2.

37. R Core Team (2020). R: A language and environment for statistical computing. R Foundation for Statistical Computing, Vienna, Austria. URL: https://www.R-project.org/.

38. Benjamini Y, Hochberg Y. Controlling the False Discovery Rate: A Practical and Powerful Approach to Multiple Testing. Journal of the Royal Statistical Society: Series B (Methodological). 1995;57:289–300. doi:10.1111/j.2517-6161.1995.tb02031.x.

