## Supplemental Figures and small tables for "Contribution of epigenetic changes to escape from X-chromosome inactivation"

### Supplemental Tables:

**Table S1: List of samples used. See additional file 1.** For CEMT samples, tissue was manually annotated to combine samples from related areas. Columns D through L are true if the dataset was available for the sample. Patient health status and sample disease are the annotations done by CEMT. CREST samples were only used for the epigenetic predictor and only samples with all datasets available were included here.

**Table S2: Comparison of histone marks between sex and XCI status. See additional file 2.** The first two sheets show BH adjusted p-values comparing female vs male and escape genes vs those subject to XCI per mark in in CEMT with our meta-status and Xi/Xa expression based XCI status calls, along with CREST data with meta-status calls and CREST data at enhancers with meta-status calls of linked genes. The last 2 sheets show the median value per mark with each sex and XCI status on the left and on the right shows the Xi/Xa ratio and log2 fold change per mark calculated based off of that median.

| | H3K4me1 | H3K4me3 | H3K9me3 | H3K27ac | H3K27me3 | ENCODE H3K27me3 | H3K36me3 | DNAme |
|---|---|---|---|---|---|---|---|---|
| **escapes from XCI** | 0.45 | 0.56 | 0.47 | 0.08 | **0.97** | **0.72** | 0.46 | 0.26 |
| **no previous call** | 0.28 | 0.15 | 0.66 | 0.01 | **0.87** | 0.49 | 0.30 | 0.29 |
| **PAR** | 0 | 0.00 | 0.28 | 0.00 | 0.04 | 0.11 | 0.17 | 0.04 |
| **subject to XCI** | 0.10 | 0.07 | **0.88** | 0.01 | **0.90** | **0.83** | 0.32 | **0.80** |
| **variably escapes from XCI** | 0.15 | 0.09 | **0.80** | 0.01 | **0.93** | **0.93** | 0.30 | 0.54 |
| **chr7** | 0.15 | 0.03 | 0.05 | 0.08 | 0.10 | 0.17 | 0.07 | 0.30 |

**Table S3: The ratio of TSSs with significant differences between males and females for various epigenetic marks using CEMT data.** The denominator was the total number of informative TSSs for which we had data. For most marks this was measured as 500bp upstream of the promoter, but for H3K36me3 we measured the mark across exons. For H3K36me3 we used unique transcripts instead of unique TSSs. Marks significant in over 70% of informative TSSs are in bold. All of the H3K27me3 data from ENCODE was downloaded and used as a replication dataset. Chromosome 7 (chr7) was included as an example autosome.

**Table S4: All XCI status calls made here. See additional file 3.** The first sheet contains a single XCI status call per gene per method. Published calls are from Balaton, *et al*. 2015. Other sheets contain all calls per sample for each method. Each row is one entry into the model, so Xi/Xa is per gene and the others are for unique transcripts. For DNAme, the samples on the far right in shades of grey are males while the samples on the left in color are females. For the epigenetic predictor, separate low confidence categories were made for when transcripts have only 12-14 of the 20 models per sample predicted a certain XCI status.

| gene | H3K4me1 | H3K4me3 | H3K9me3 | H3K27ac | H3K27me3 | H3K36me3 | DNAme | expression | nE | nS |
|---|---|---|---|---|---|---|---|---|---|---|
| BCOR | 0.076 | 0.42 | **0.019** | 0.43 | 0.065 | 0.56 | **0.011** | **0.0097** | 2 | 5 |
| CXorf38 | 0.071 | 0.26 | 0.82 | 0.52 | 0.18 | 0.24 | 0.10 | 0.18 | 3 | 2 |
| EIF2S3 | 0.054 | 0.097 | 0.80 | 0.12 | 0.33 | 0.54 | **0.040** | 0.070 | 4 | 2 |
| MED14 | 0.66 | 0.98 | 0.32 | 0.39 | 0.28 | 0.45 | 0.42 | 0.86 | 2 | 3 |
| PNPLA4 | 0.84 | **0.029** | **0.0076** | 0.069 | 0.070 | 0.42 | 0.15 | 0.74 | 4 | 2 |
| PRKX | 0.73 | 0.070 | **0.029** | **0.047** | 0.27 | **0.021** | **0.048** | 0.053 | 5 | 3 |
| SMC1A | 0.11 | **0.043** | 0.24 | 0.61 | **0.036** | **0.048** | **0.046** | 0.15 | 6 | 2 |
| TIMP1 | 0.59 | 0.40 | 0.55 | 0.054 | 0.78 | 0.18 | 0.21 | 0.87 | 3 | 4 |

**Table S5: Significance of differences in epigenetic marks between samples with opposite XCI statuses at genes found variably escaping XCI by Xi/Xa expression.** We also tested whether expression differed between samples with opposite XCI status. Presented here are the p-values of t-tests. Those with p-values less than 0.05 are in bold. nE and nS are the number of samples escaping or subject to XCI for each gene.

| gene | H3K4me1 | H3K4me3 | H3K9me3 | H3K27ac | H3K27me3 | H3K36me3 | DNAme | expression |
|---|---|---|---|---|---|---|---|---|
| PRKX | 2.207482 | 10.81617 | **-2.32661** | **4.171272** | -2.94785 | **1.586497** | **-4.70569** | 0.460988 |
| PNPLA4 | 1.159099 | **10.4815** | -14.1268 | 8.604572 | -3.90181 | -1.2279 | -17.5906 | 0.887235 |
| PNPLA4 | 1.088554 | 6.106866 | **-7.22367** | 3.873968 | -0.5687 | -1.2279 | -13.0625 | 0.79953 |
| PNPLA4 | 1.088554 | 6.106866 | **-7.22367** | 3.873968 | -0.5687 | -0.50595 | -13.0625 | 1.070425 |
| EIF2S3 | 4.328741 | 12.59848 | 0.666695 | -7.88552 | -2.50986 | 0.42919 | **-30.1127** | 56.54015 |
| BCOR | 2.831612 | -1.35245 | -0.25747 | 1.548647 | 1.395386 | -0.26325 | 15.31005 | **32.60517** |
| BCOR | 5.870941 | 1.363537 | **-0.54052** | 1.472802 | -5.61021 | -0.26325 | **-22.549** | **33.10998** |
| BCOR | 5.870941 | 1.363537 | **-0.54052** | 1.472802 | -5.61021 | 0.016699 | **-22.549** | **32.64655** |
| CXorf38 | 13.65249 | 3.909036 | -1.12366 | 1.58118 | -3.6874 | -0.11738 | -54.6914 | 10.16366 |
| CXorf38 | 16.67418 | 2.714651 | -0.85511 | 1.846735 | -4.25266 | -0.35507 | -36.7491 | 10.28184 |
| MED14 | 1.039564 | 3.233193 | -1.08359 | 0.868199 | -3.15195 | -0.18656 | -0.58621 | 0.365396 |
| TIMP1 | 3.746742 | -0.83792 | 0.437359 | -0.83214 | -1.3243 | -0.34116 | 14.47131 | 0.016658 |
| SMC1A | -8.21266 | **8.614903** | -9.708 | 1.579612 | **-5.34306** | **-0.22771** | **-43.8589** | -0.06352 |
| SMC1A | -8.21266 | **8.614903** | -9.708 | 1.579612 | **-5.34306** | -0.1797 | **-43.8589** | -0.08209 |

**Table S6: The differences in epigenetic marks between samples with opposite XCI statuses at genes found variably escaping XCI by Xi/Xa expression.** The mean value for samples subject to XCI was subtracted from the mean value for samples escaping XCI. Those found significant in Table 1 are bolded. Genes with multiple transcripts are included multiple times, even if they share a TSS.

| | |
|---|---|
| **H3K4me1** | 0.093906 |
| **H3K4me3** | **3.34E-11** |
| **H3K9me3** | **7.66E-68** |
| **H3K27ac** | **8.11E-09** |
| **H3K27me3** | **1.03E-29** |
| **H3K36me3** | 0.067493 |

**Table S7: adjusted p-values comparing marks in females between genes found subject to XCI vs escaping XCI by DNAme.** Those significant are in bold (adjusted p-value<0.01).

| **bin** | **<25%** | 33-66% | >75% |
|---|---|---|---|
| **0** | 0.990521 | 0 | 0 |
| **0-10%** | 0.967776 | 0.011134 | 0.002227 |
| **10-20%** | 0.731334 | 0.138481 | 0.035099 |
| **20-30%** | 0.591919 | 0.172437 | 0.130438 |
| **30-40%** | 0.521443 | 0.128352 | 0.242815 |
| **40-50%** | 0.447618 | 0.093807 | 0.359666 |
| **50-60%** | 0.337163 | 0.11472 | 0.44407 |
| **60-70%** | 0.213763 | 0.151308 | 0.5045 |
| **70-80%** | 0.08201 | 0.16705 | 0.602591 |
| **80-90%** | 0.026355 | 0.094393 | 0.755547 |
| **90-100%** | 0.0059 | 0.02496 | 0.914908 |

**Table S8: Distribution summary for DNAme per read.** The number is what percent of reads in each bin were below 25%, between 33 and 66% or over 75% DNAme.

| Histone Mark | Accuracy for genes escaping XCI | Accuracy for genes subject to XCI |
|---|---|---|
| **H3K4me1** | 0.27 | 0.45 |
| **H3K4me3** | 0.36 | 0.55 |
| **H3K9me3** | 0.11 | 0.16 |
| **H3K27ac** | 0.33 | 0.64 |
| **H3K27me3** | 0.51 | 0.21 |
| **H3K36me3** | 0.09 | 0.03 |

**Table S9: The accuracy of simple models predicting XCI status from a single histone mark.** These accuracies are low because the models overpredicted variable escape from XCI as there is large overlap between the two XCI statuses.

| Histone Mark | Accuracy for genes escaping XCI | Accuracy for genes subject to XCI |
|---|---|---|
| **H3K4me1** | 0.5055 | 0.6262 |
| **H3K4me3** | 0.6196 | 0.7497 |
| **H3K9me3** | 0.6097 | 0.7015 |
| **H3K27ac** | 0.5405 | 0.6888 |
| **H3K27me3** | 0.6609 | 0.7383 |
| **H3K36me3** | 0.5333 | 0.6115 |
| **DNAme** | 0.826 | 0.8816 |

**Table S10: The accuracy of random forest models predicting XCI status from a single histone mark.**

| | No CpG island | CpG island | low expression | high expression |
|---|---|---|---|---|
| escapes XCI | 8 | 95 | 26 | 77 |
| subject to XCI | 600 | 1116 | 549 | 1167 |
| variably escapes XCI | 2 | 7 | 1 | 8 |
| inconsistent prediction | 462 | 346 | 569 | 239 |

**Table S11: XCI status calls made using a random forest epigenetic predictor with all of the epigenetic marks, split by presence or absence of a CpG island and expression.** The threshold used to split low from high expression is a median of 0.1 RPKM across samples. Inconsistent predictions had over a third of samples with fewer than 15 of the 20 models trained agree on an XCI status.

| | Variable escape across individuals | Variable escape across tissues | | Variable escape across TSSs | |
|---|---|---|---|---|---|
| Total number of genes | 9 | 65 | 2636 | 1 | 461 |
| H3K27me3 TSS | 0 | **51%** | 27% | 0 | 12% |
| H3K27me3 genebody | 0 | 28% | 41% | 0 | 22% |
| H3K27ac TSS | 0 | 31% | 37% | 100% | 17% |
| H3K9me3 TSS | 0 | 32% | 19% | 0 | 30% |
| H3K9me3 genebody | 11% | 8% | 7% | 100% | 29% |
| H3K4me3 TSS | 0 | **40%** | 20% | 0 | 22% |
| H3K4me1 TSS | 0 | 62% | 53% | 100% | 19% |
| H3K36me3 TSS | 0 | 35% | 22% | 0 | 12% |
| H3K36me3 genebody | 11% | 29% | 31% | 0 | 38% |
| DNAme TSS | 67% | **58%** | 28% | 100% | 19% |
| expression | 0 | **38%** | 1% | | |

**Table S12: The percent of genes found variably escaping by our epigenetic predictor with significant differences in various epigenetic marks.** Genes were counted as significant if BH corrected p-values were less than 0.01 when comparing samples predicted as subject to XCI to samples predicted as escaping from XCI. The total number of genes row shows the total number of genes in each category. The variable escape across tissues and TSSs categories have 2 columns each, the left column being the percent of variably escaping genes with significant differences between tissues/TSSs and the right column being the percent of all genes on the X chromosome with differences between tissues/TSSs. Highlighted in blue are marks which were significantly more likely to have significant differences between tissues/TSSs at genes predicted to variably escape than in all X linked genes.

| variable escape threshold | 33% | 25% | 10% | 5% |
|---|---|---|---|---|
| Number of genes variably escaping across samples | 9 | 41 | 431 | 740 |
| H3K27me3 TSS | 0% | 4.90% | 23% | 27% |
| H3K27me3 genebody | 0% | 0% | 3.70% | 4.20% |
| H3K27ac TSS | 0% | 2% | 3.50% | 7.80% |
| H3K9me3 TSS | 0% | 0% | 11% | 15% |
| H3K9me3 genebody | 11% | 0% | 2.10% | 4.30% |
| H3K4me3 TSS | 0% | 0% | 2.80% | 4.20% |
| H3K4me1 TSS | 0% | 4.90% | 7.70% | 10% |
| H3K36me3 TSS | 0% | 2.40% | 13% | 16% |
| H3K36me3 genebody | 11% | 7% | 3.50% | 6.20% |
| DNAme | 67% | 22% | 17% | 20% |
| expression | 0% | 0% | 0.90% | 0.70% |

**Table S13: The percent of genes found variably escaping by our epigenetic predictor with significant differences in various epigenetic marks across various variable escape thresholds.** Variable escape threshold is the number of samples with each XCI status (escaping from XCI and subject to XCI) that were required in order to call a gene as variably escaping from XCI across samples. Genes were counted as significant if BH corrected p-values were less than 0.01 when comparing samples predicted as subject to XCI to samples predicted as escaping from XCI.

| | CEMT calls per gene | | | |
|---|---|---|---|---|
| CREST calls per gene | | Escapes XCI | Subject to XCI | Variably escapes XCI |
| | Escapes XCI | 64 | 37 | 7 |
| | Subject to XCI | 3 | 1492 | 1 |
| | Variably escapes XCI | 1 | 32 | 0 |

**Table S14: Comparing XCI status calls made by an epigenetic predictor in the CEMT dataset vs a similar model in the CREST dataset.**

| | VE across individuals | VE across tissues | | VE across TSSs | |
|---|---|---|---|---|---|
| **total** | **53** | **13** | **2313** | **6** | **1155** |
| **H3K4me1.Fval** | 0 | 0 | 0.076% | 0 | 0.087% |
| **H3K4me1.Mval** | 0 | <NA> | NA | 17% | 4.2% |
| **H3K4me3.Fval** | 0 | 0 | 0 | **50%** | **1%** |
| **H3K4me3.Mval** | 0 | <NA> | NA | **67%** | **6%** |
| **H3K9me3.Fval** | 0 | 0 | 0 | 0 | 0 |
| **H3K9me3.Mval** | 0 | <NA> | NA | 33% | 5% |
| **H3K27Ac.Fval** | 0 | 0 | 0.08% | 17% | 1% |
| **H3K27Ac.Mval** | 0 | <NA> | NA | **50%** | **5%** |
| **H3K27me3.Fval** | 0 | **15%** | **0.11%** | **17%** | **9%** |
| **H3K27me3.Mval** | 0 | <NA> | NA | **33%** | **2%** |
| **H3K36me3.Fval** | 0 | 0 | 0 | 0 | 2% |
| **H3K36me3.Mval** | 0 | <NA> | NA | **50%** | **3%** |
| **WGBS.Fval** | 2% | **92%** | **2.31%** | **100%** | **6%** |
| **WGBS.Mval** | 0 | <NA> | NA | **100%** | **1%** |
| **RNAseq.Fval** | 4% | **46%** | **1.33%** | 0 | 3% |
| **RNAseq.Mval** | 0 | <NA> | NA | 17% | 3% |

**Table S15: The percent of genes found variably escaping by our epigenetic predictor in the CREST dataset with significant differences in various epigenetic marks.** Genes were counted as significant if BH corrected p-values were less than 0.01 when comparing samples predicted as subject to XCI to samples predicted as escaping from XCI. The total number of genes row shows the total number of genes in each category. The variable escape across tissues and TSSs categories have 2 columns each, the left column being the percent of variably escaping genes with significant differences between tissues/TSSs and the right column being the percent of all genes on the X chromosome with differences between tissues/TSSs. Highlighted in blue are marks which were significantly more likely to have significant differences between tissues/TSSs at genes predicted to variably escape than in all X linked genes.

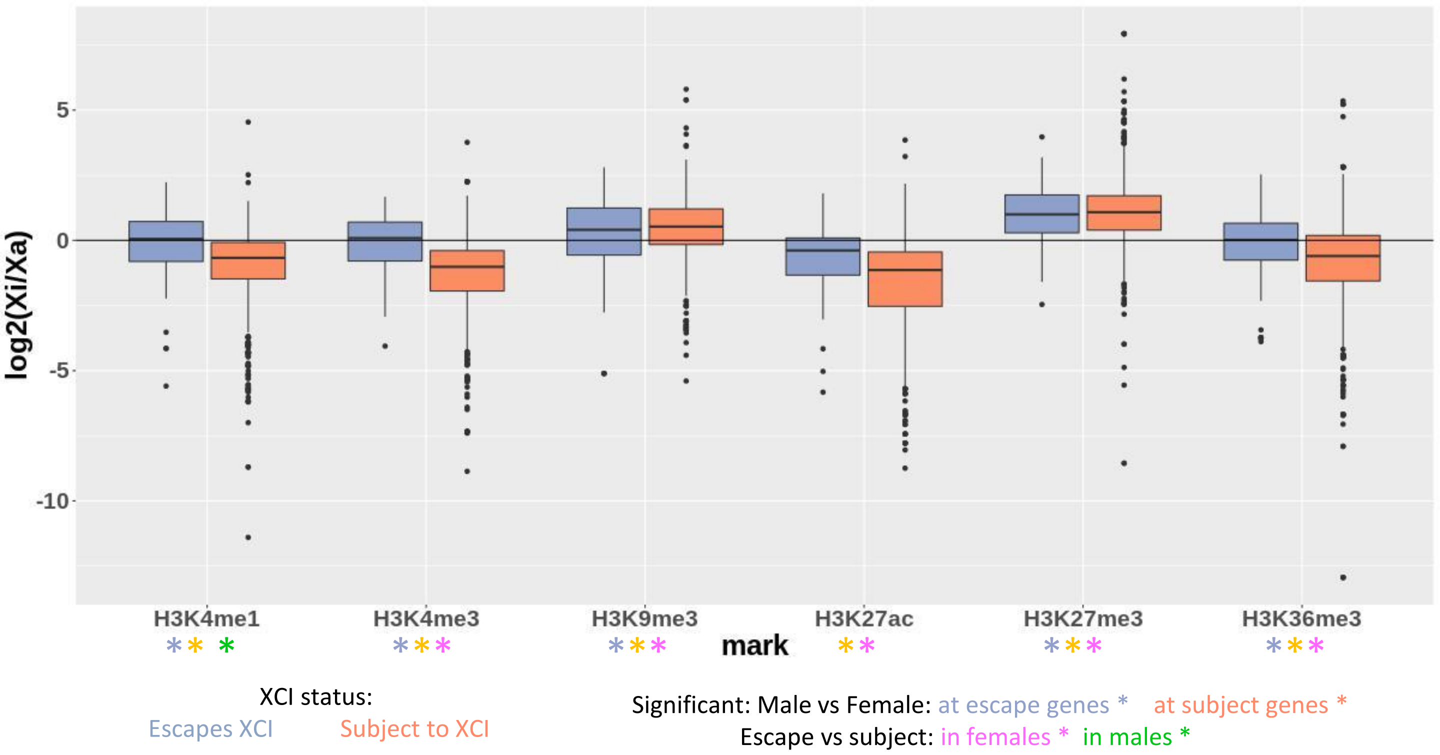


**Figure S1: log2(Xi/Xa) for epigenetic marks in CREST at promoters.** Data from CREST is shown. Significance for the various t-tests featured in Table S2 are shown by the differently colored star.

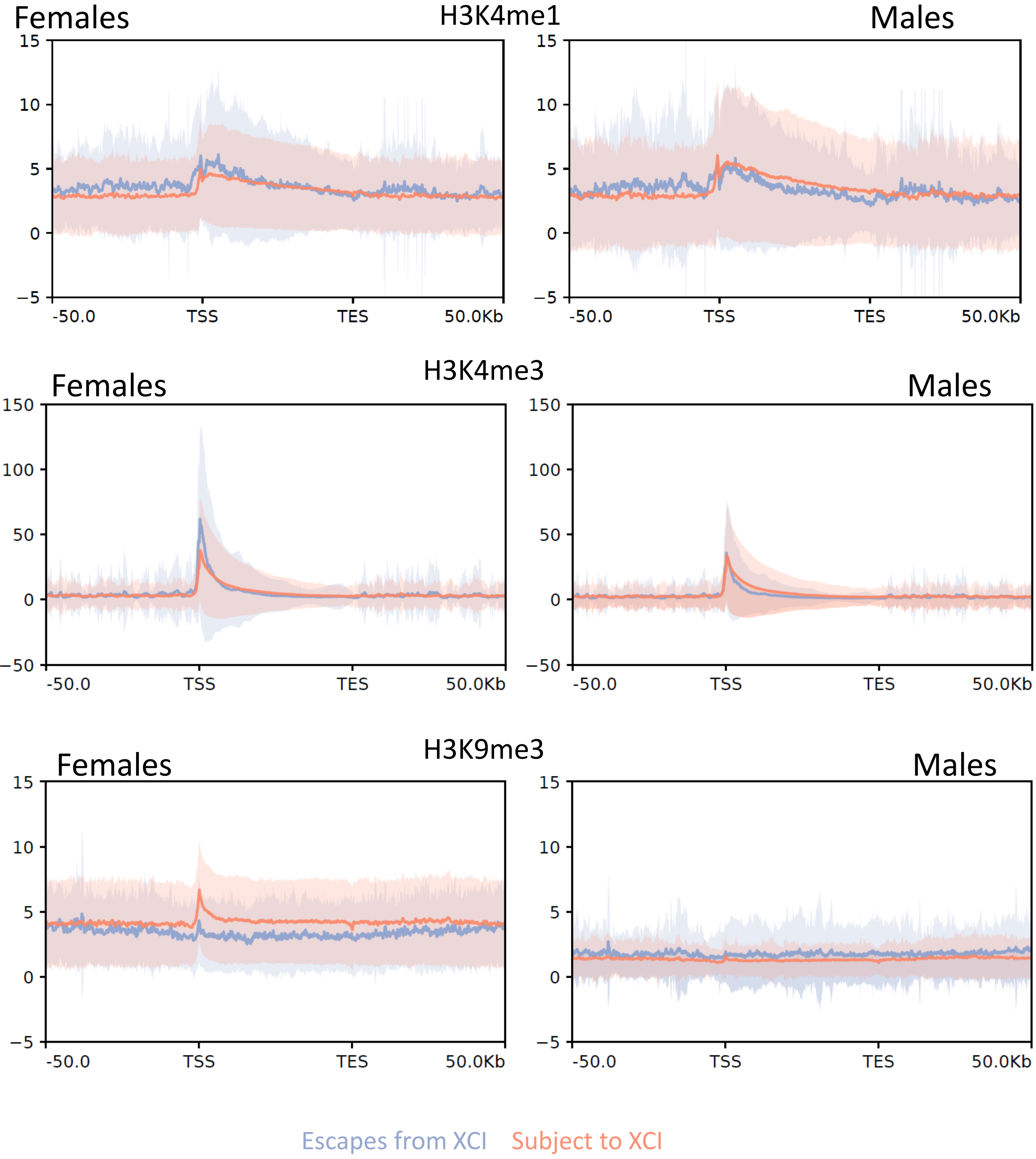


**Figure S2: Meta-gene plots of histone marks within 50kb of genes, separated by XCI status.** The plots were generated using deeptools computeMatrix and plotProfile on bigwig files that were the mean across samples. Lighter shaded regions show the standard deviation of each mark. Continued on next page

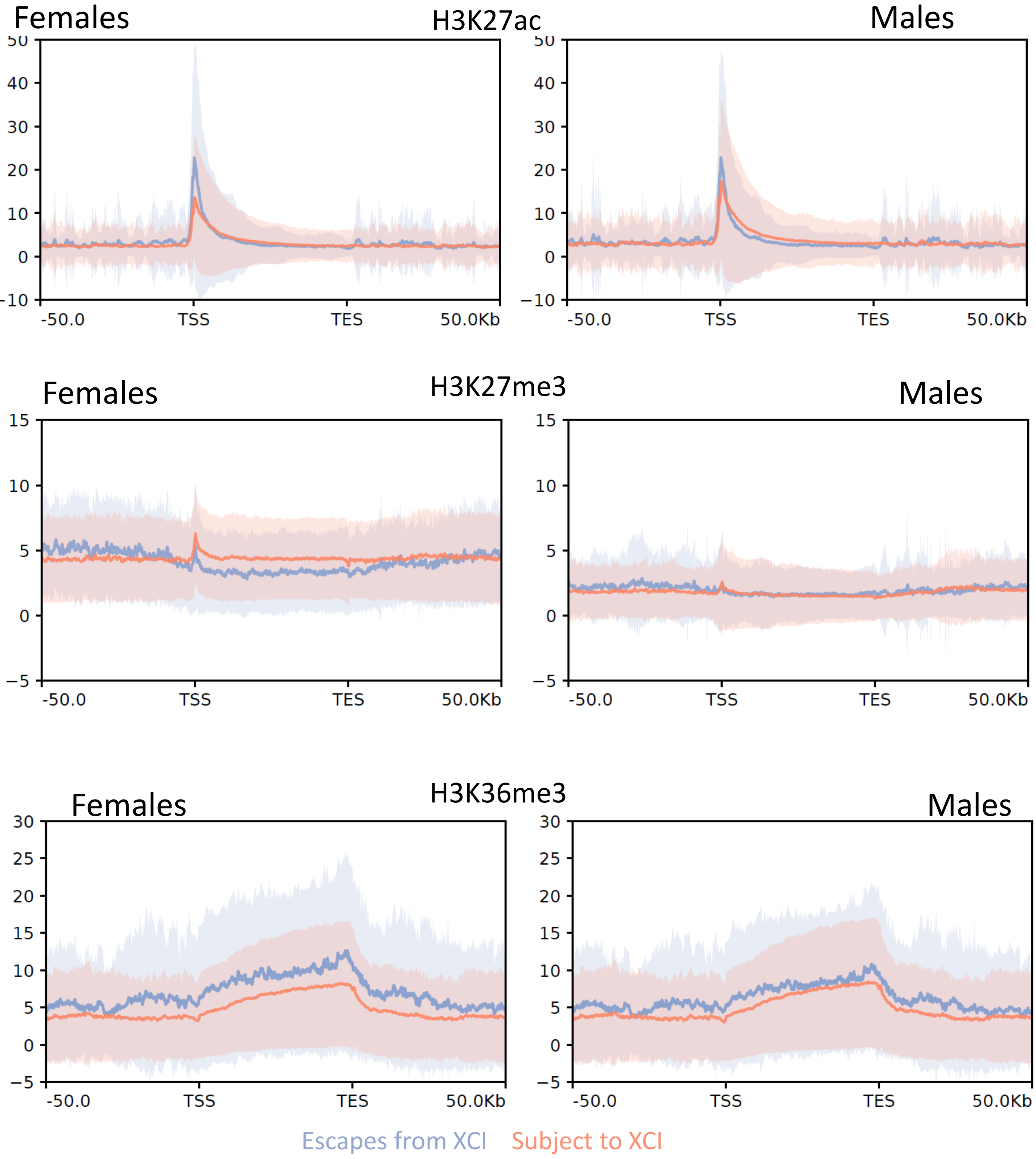


**Figure S2: Continued from previous page**

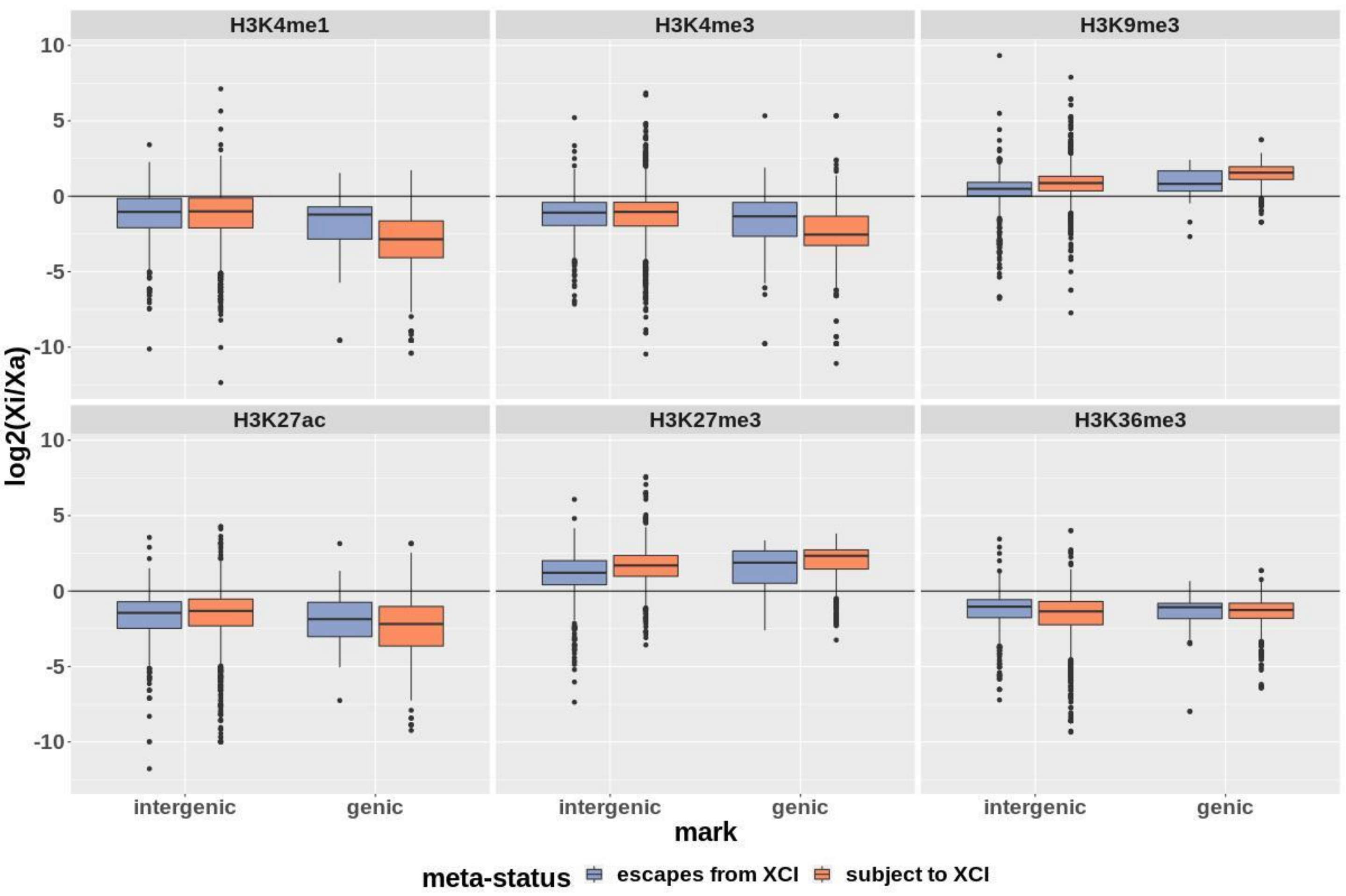


**Figure S3: log2(Xi/Xa) for epigenetic marks in CEMT at enhancers mapping to genes which escape from or are subject to XCI.** Enhancers are split by whether they are located within a gene (genic) or not (intergenic).

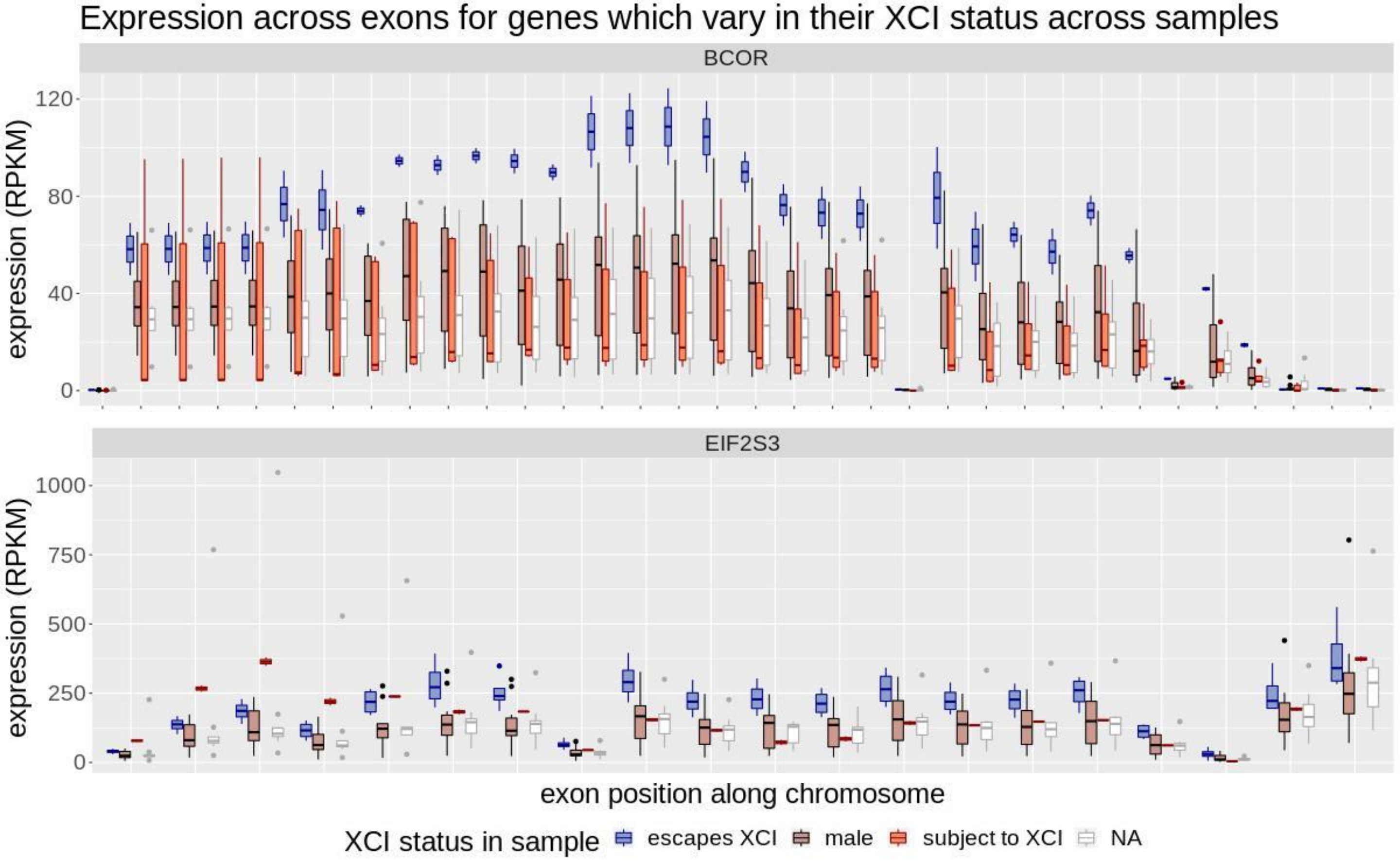


**Figure S4: Expression across exons for genes with significantly different expression in samples with opposite XCI statuses.** XCI status per sample was determine here using Xi/Xa expression.

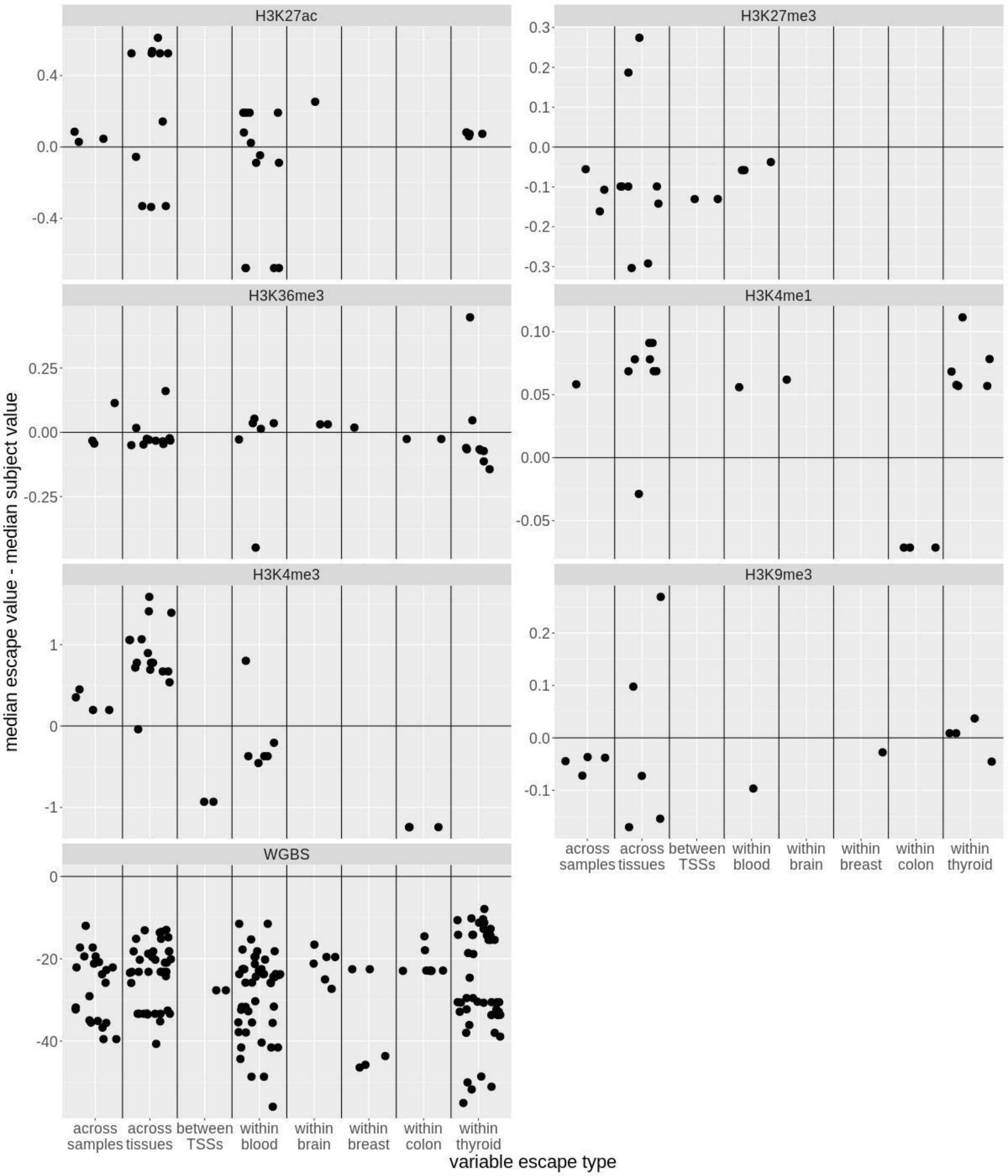


**Figure S5: Differences in epigenetic marks between samples found escaping vs subject to XCI at variably escaping genes in DNAme.** For most of these marks, the region 500bp upstream of the promoter is used, except for H3K36me3 which uses the gene body.

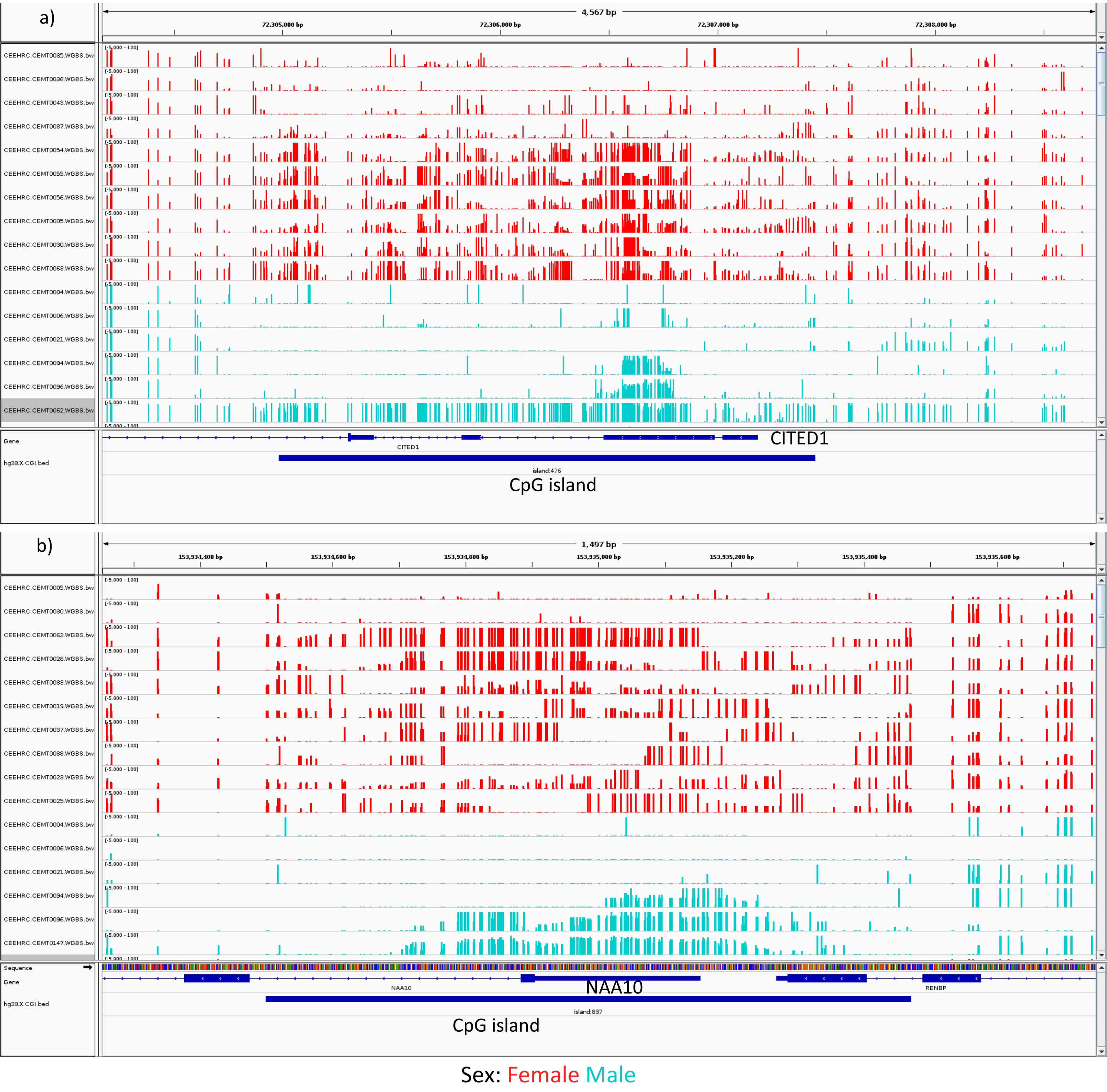


**Figure S6: IGV view of DNAme bigwig tracks at two variably escaping genes.** a) A view of the CpG island at CITED1. b) a view of the CpG island at NAA10. A broad representation of samples was sought, some hypomethylated, some hypermethylated and some inconsistent across the CpG island. Broad hypermethylation in males at these genes was rare but is included here as an example of an extreme.

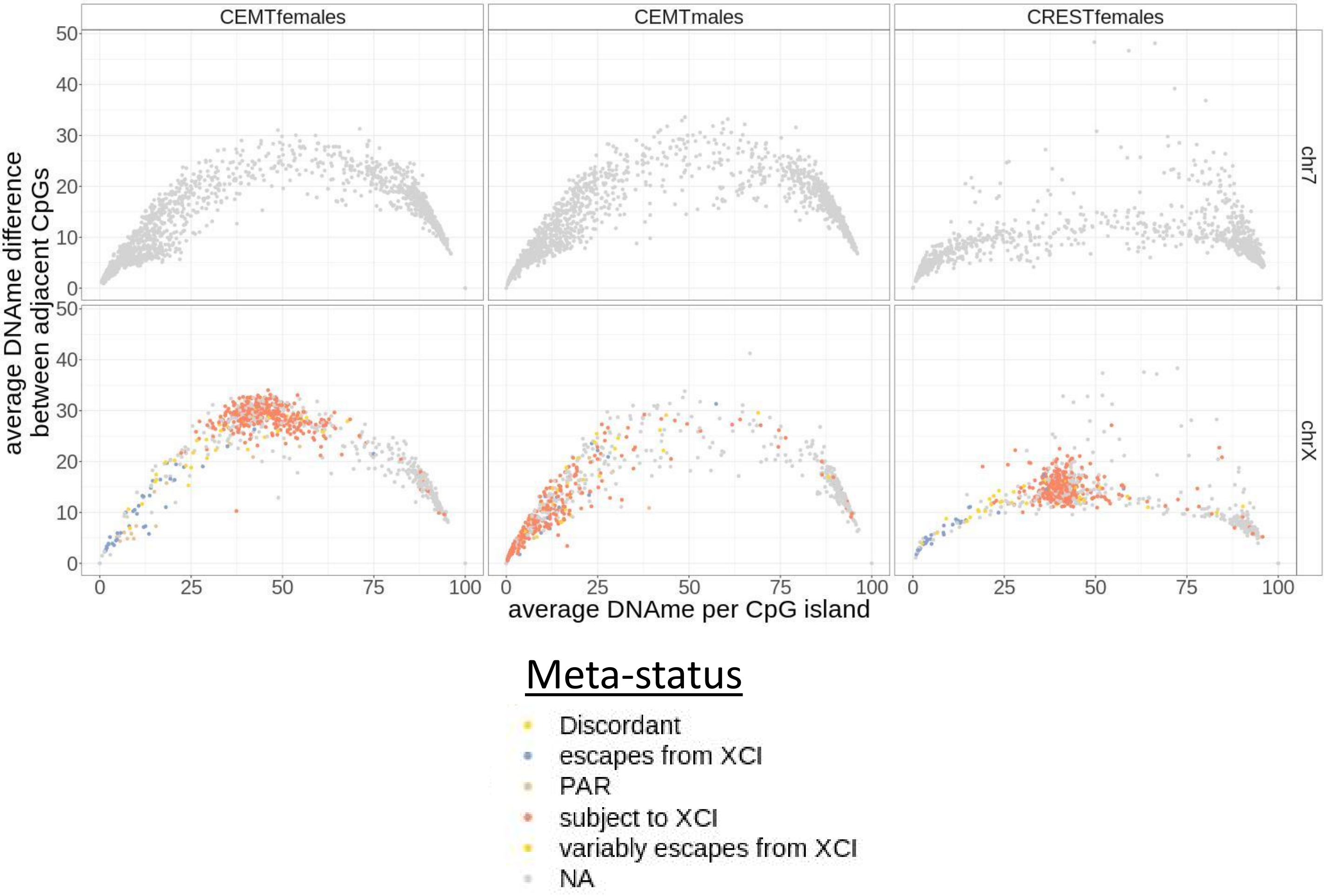


**Figure S7: average DNAme difference between adjacent CpGs per CpG island.** Each point is the average DNAme difference between adjacent CpGs for an individual island, averaged again across samples. Islands are colored by the meta-status of the closest TSS within 2kb. Chr7 was chosen as an autosomal control to show whether the differences are X specific. Males and females from CEMT were used to check for sex specificity and females from CREST were included to check for cancer specificity.

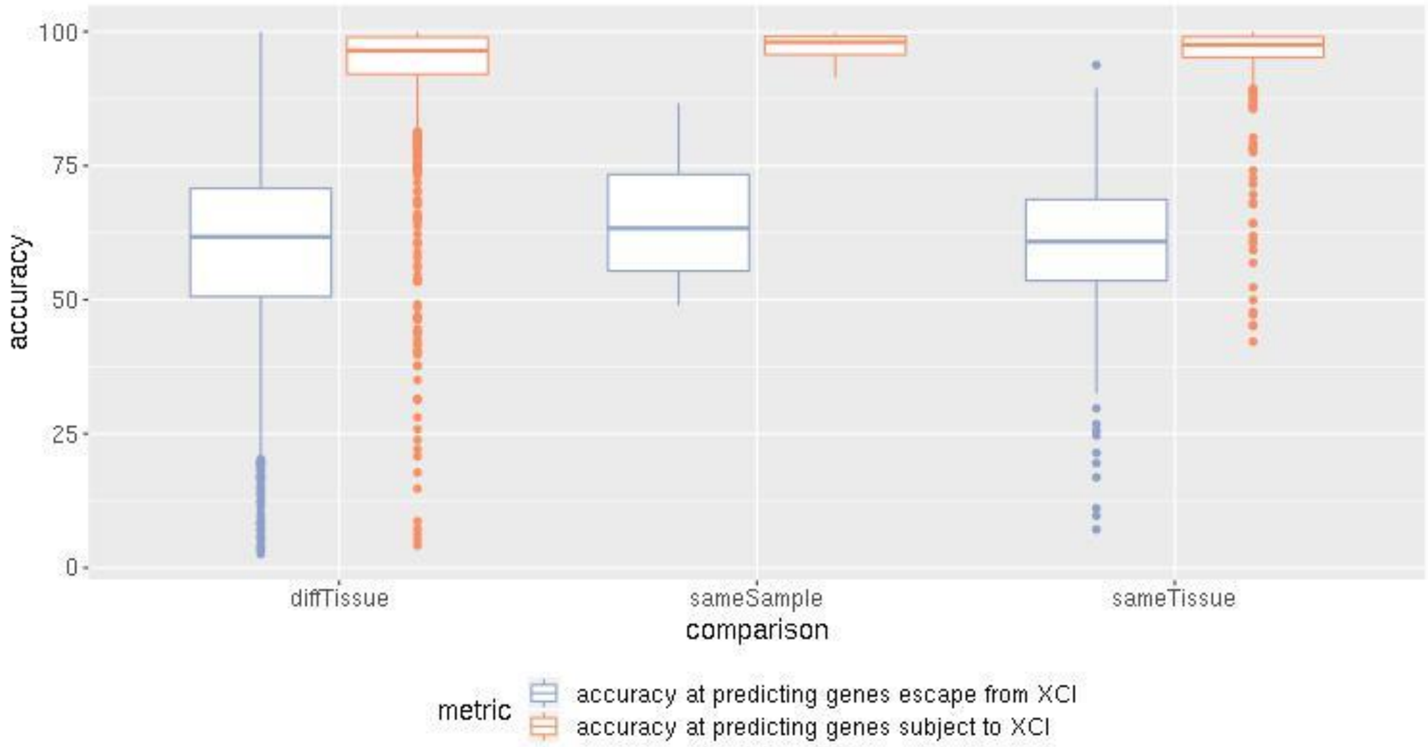


**Figure S8: Accuracy when models trained in one sample are tested on other models.**

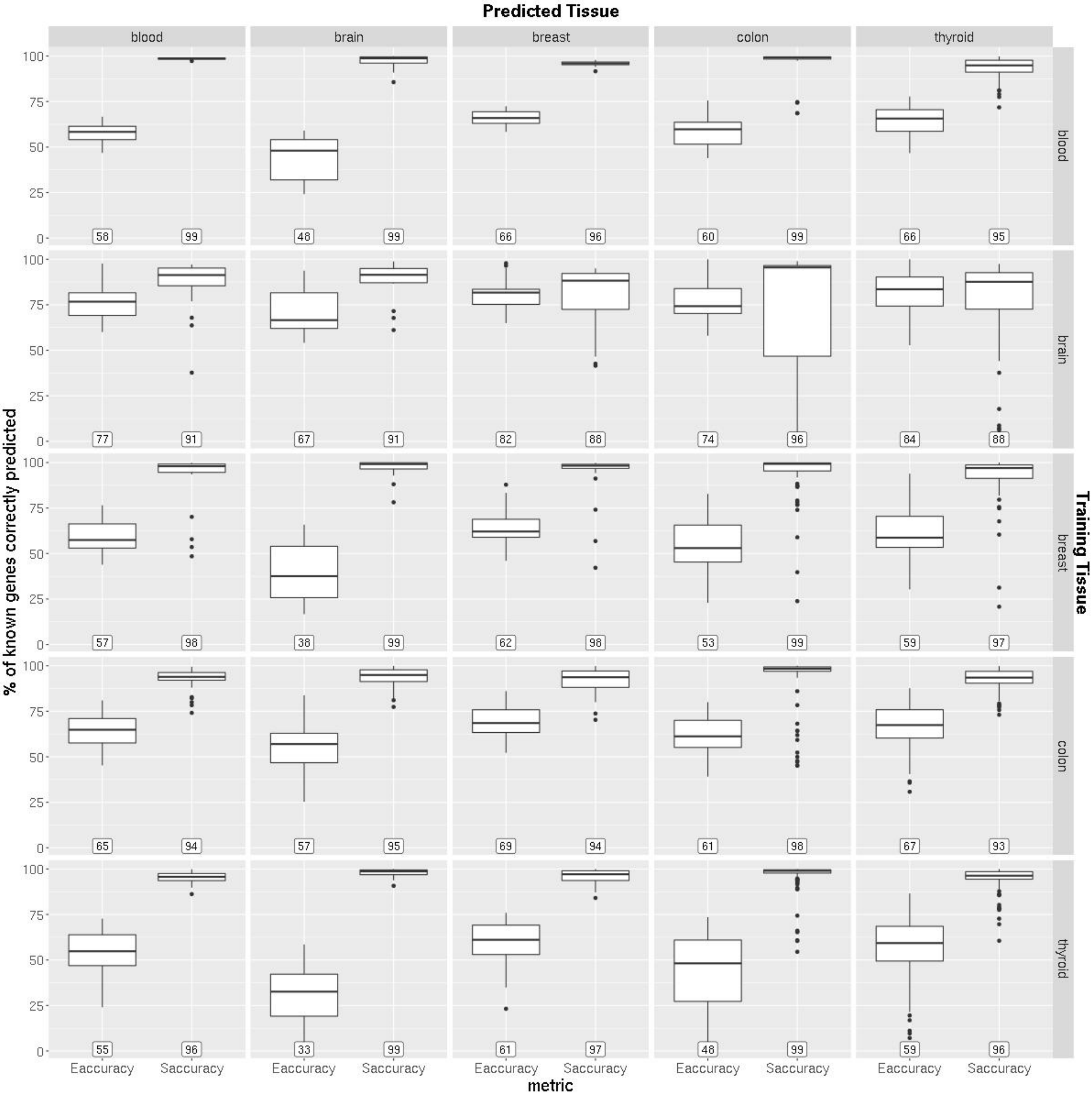


**Figure S9: Accuracy when models trained in one sample are tested on other models, separated per tissue comparison.** The numbers at the bottom of each plot are the median accuracy. Each point is the accuracy at predicted an XCI status when a model from the training tissue on a sample in the predicted tissue. Eaccuracy is accuracy at predicting genes as escape from XCI. Saccuracy is accuracy when predicting genes as subject to XCI.

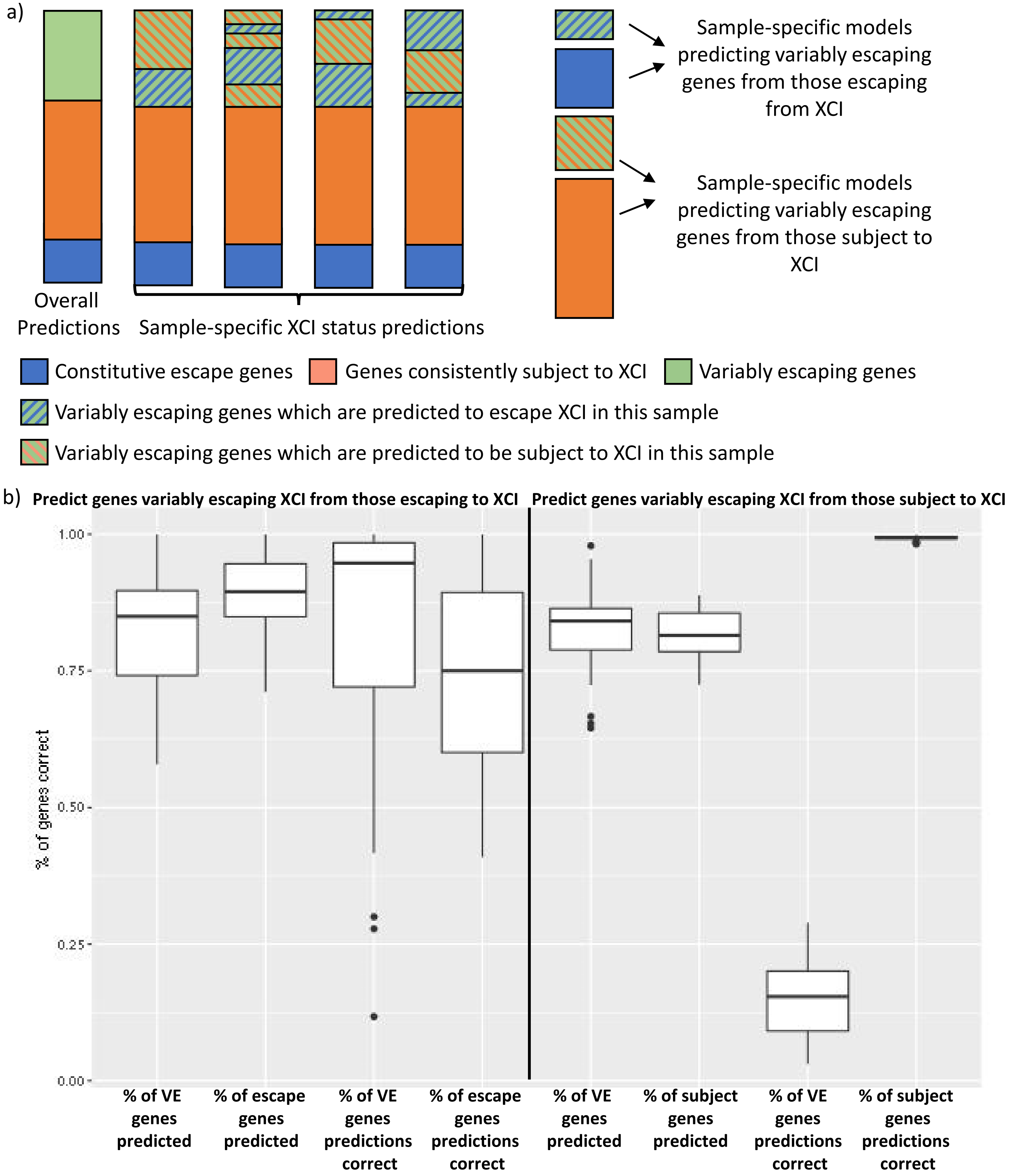


**Figure S10: Models to predict variably escaping genes from those with a consistent XCI status.** (a) a diagram depicting our models to predict genes which are variably escaping from those consistently escaping or subject to XCI. (b) Accuracy metrics when predicting which genes variably escape from XCI across samples using data from individual samples. On the left are metric when a model is trained on only genes called as escaping XCI in that sample, while the right is metrics when a model is trained on only genes called as subject to XCI. VE is variably escaping from XCI. Subject is subject to XCI.

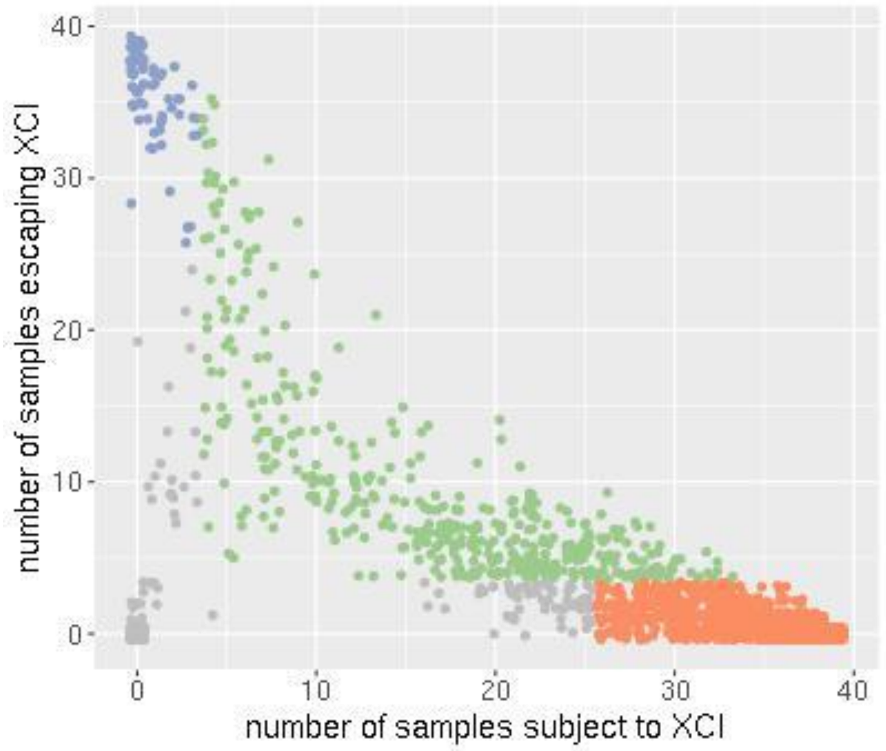


**Predicted XCI Status:**

Escapes XCI   Subject to XCI   Variably escapes from XCI   unknown

**Figure S11: The number of samples called as escaping vs subject to XCI per transcript by our epigenetic predictor.**

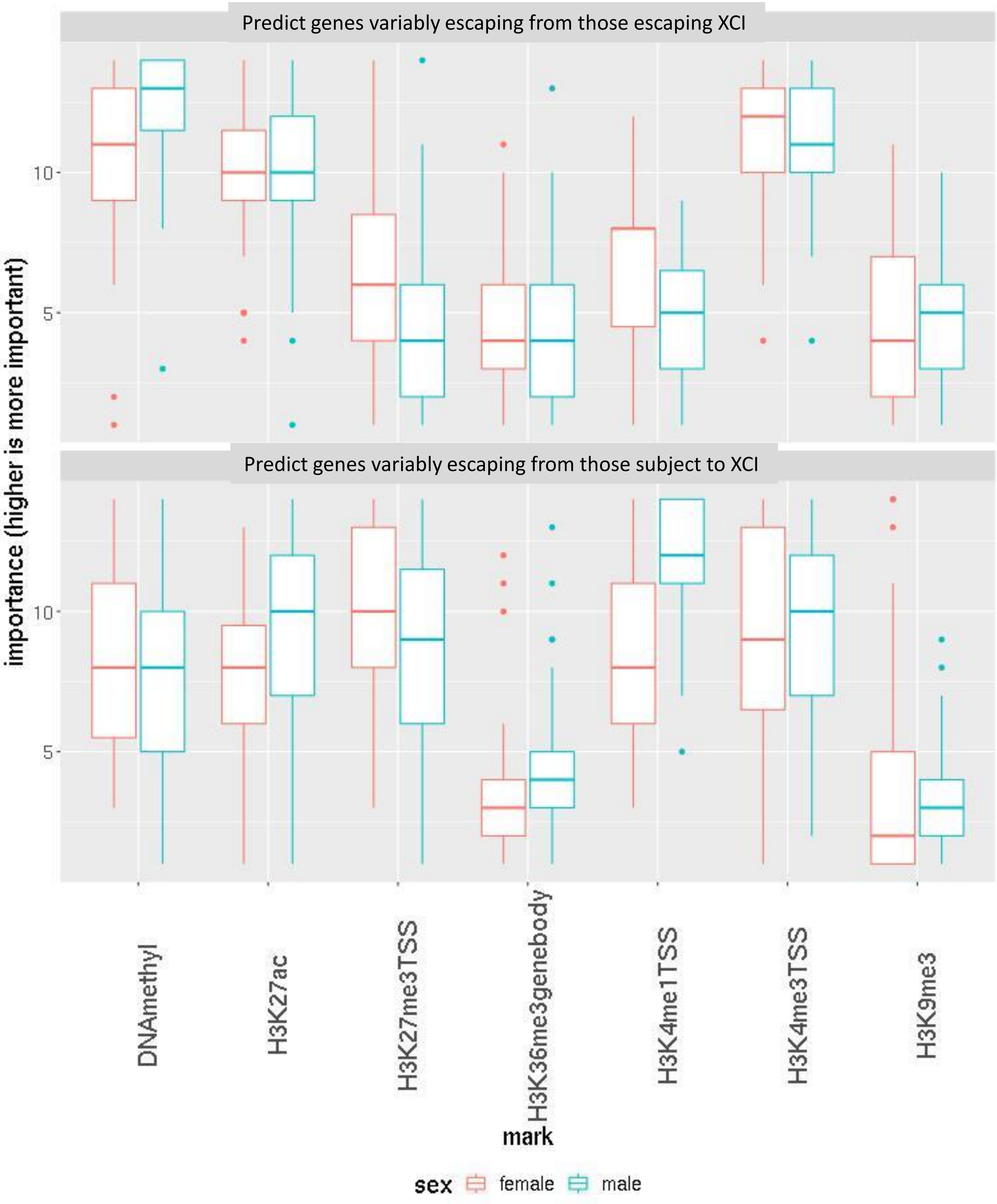


**Figure S12: Ranked importance of the marks used to predict which genes variably escape across samples.** The contributions to each model from each mark were ranked, with rank 14 being the most important and rank one being the least important.

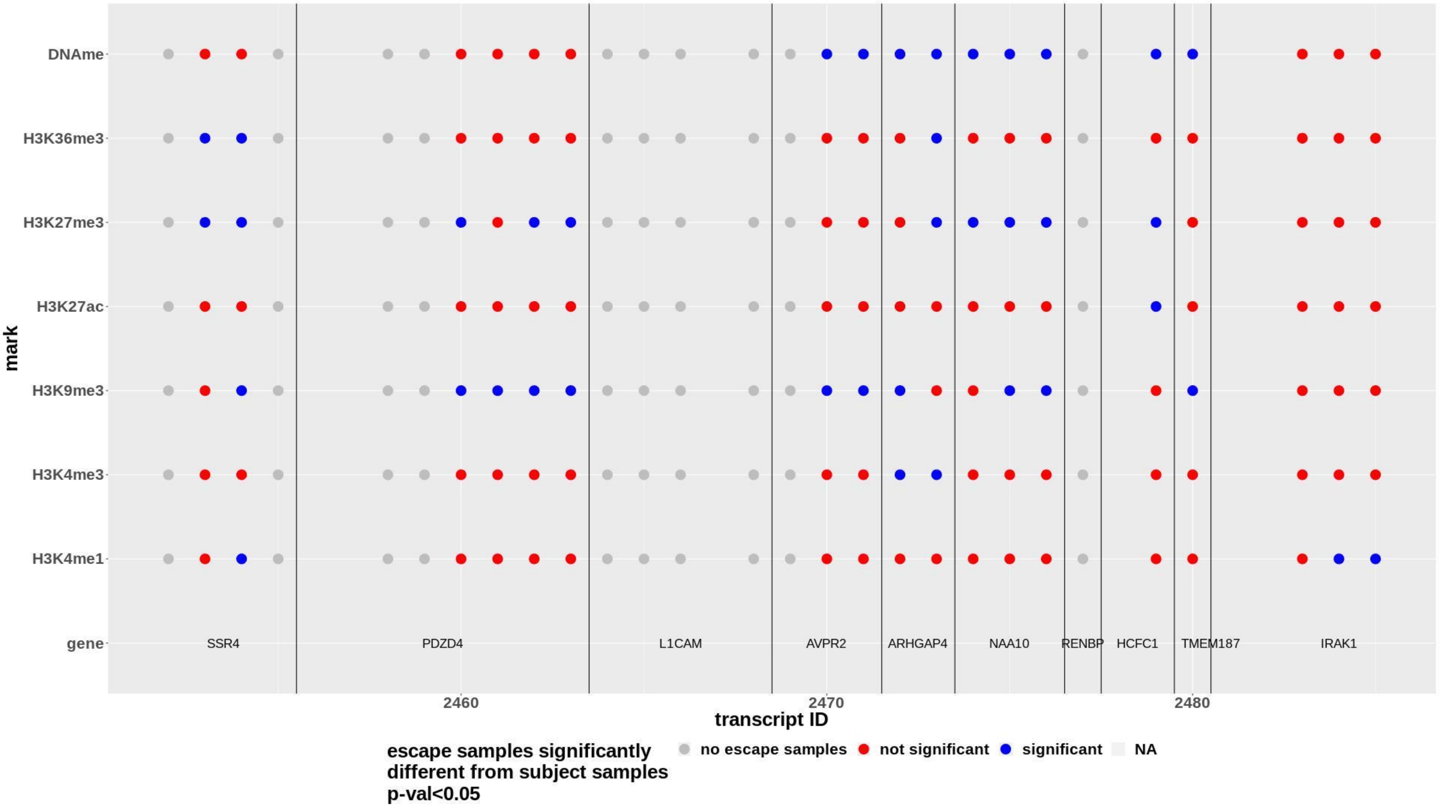


**Figure S13: Which marks were significantly different between samples predicted as escaping vs subject to XCI in a variably escaping region.** Transcript ID is the order that the transcripts are located along the chromosome. There are multiple transcripts per gene but they may be sharing the same TSS and have the same data for all marks but H3K36me3. Vertical lines are drawn denoting which transcripts belong with each gene.
